# The innate sensor ZBP1-IRF3 axis regulates cell proliferation in multiple myeloma

**DOI:** 10.1101/2020.06.17.157107

**Authors:** Kanagaraju Ponnusamy, Maria Myrsini Tzioni, Murshida Begum, Mark E Robinson, Valentina S Caputo, Alexia Katsarou, Nikolaos Trasanidis, Xiaolin Xiao, Ioannis V Kostopoulos, Deena Iskander, Irene Roberts, Pritesh Trivedi, Holger W Auner, Kikkeri Naresh, Aristeidis Chaidos, Anastasios Karadimitris

**Affiliations:** Hugh & Josseline Langmuir Centre for Myeloma Research, Centre for Haematology, Department of Immunology & Inflammation, Imperial College London, London, United Kingdom; Department of Haematology, Hammersmith Hospital, Imperial College Healthcare NHS Foundation Trust, London, United Kingdom; Section of Animal and Human Physiology, National and Kapodestrian University of Athens, Department of Biology, School of Science, Athens, Greece; Department of Paediatrics and MRC Molecular Haematology Unit, Weatherall Institute of Molecular Medicine, University of Oxford and BRC Blood Theme, NIHR Oxford Biomedical Centre, Oxford, United Kingdom; Department of Cellular & Molecular Pathology, Northwest London Pathology, Imperial College Healthcare NHS Trust, London, United Kingdom

**Keywords:** ZBP1, multiple myeloma, IRF3, IRF4, TBK1, cell cycle

## Abstract

ZBP1 is an inducible, non-constitutively expressed cellular nucleic acid sensor that triggers type I interferon (IFN) responses via phosphorylation and activation of the transcription factor (TF) IRF3 by TBK1. However, the role of the ZBP1-IRF3 axis in cancer is not known. Here we show that ZBP1 is selectively and constitutively expressed in late B cell development and it is required for optimal T cell-dependent humoral immune responses. In the plasma cell (PC) cancer multiple myeloma, interaction of constitutively expressed ZBP1 with TBK1 and IRF3 results in IRF3 phosphorylation. Notably, rather than IFN type I response genes, IRF3 directly activates, in part through co-operation with the PC lineage-defining TF IRF4, cell cycle genes thus promoting myeloma cell proliferation. This generates a novel, potentially therapeutically targetable and relatively selective myeloma cell addiction to the ZBP1-IRF3 axis. These data expand our knowledge of the role of cellular immune sensors in cancer biology.

---

ZBP1 is one of a group of cellular DNA/RNA sensors that activate innate immunity in response to pathogen-derived or cellular nucleic acids (NA)^1–3^. Binding of double stranded Z-form RNA (dsRNA) to its Zα and Zβ domains is followed by a RHIM-RHIM domain interaction of ZBP1 with RIPK3 and induction of programmed cell death^4–6^. This process is antagonised by RIPK1, preventing for example, ZBP1-dependent cell death and inflammation in the developing skin^7,8^.

Alternatively, ZBP1 directly or through interaction with the cellular NA sensor cGAS-STING axis, can lead to phosphorylation of the transcription factor IRF3 (pIRF3) by the serine/threonine kinase TBK1. pIRF3 after nuclear entry acquires the ability to activate transcription of interferon (IFN) type I response genes^1,9–11^. Unlike other NA sensors, expression of ZBP1 is inducible and triggered in response to NA binding to its Zα and Zβ domains^6,12,13^.

As well as in innate immune responses, the cGAS-STING-IRF3 axis is also implicated in cancer by exerting tumor cell intrinsic or extrinsic onco-suppressive effects in response to excess cellular cytosolic DNA or to activated endogenous retroviruses in transformed cells^14–16^. In some cellular contexts, cGAS might promote oncogenesis through inhibition of DNA repair pathways^17,18^. Chronically activated cGAS–STING has been linked with persistent inflammation and cancer progression while its downstream target TBK1 has been shown to co-operate with oncogenic RAS to promote cancer cell survival^19^.

Multiple myeloma (MM) is a common incurable blood cancer of the bone marrow (BM) plasma cells (PC), the immunoglobulin-secreting terminally differentiated B lineage cells^20–22^. Primary and secondary somatic genetic events comprising copy number and single nucleotide variants shape a genomic landscape of extensive, in time and in space, genetic heterogeneity and diversification rendering targeted therapies for MM a challenging task cells^20–22^. In this regard, there is a need for identification of biological pathways that are involved in myelomagenesis independent of the genetic status and extend of diversification. A search that we undertook to address this requirement identified ZBP1 as selectively expressed in myeloma PC. Prompted by this finding, we explored the potential role of ZBP1 in humoral immune responses and dissected its function in myeloma cell survival and proliferation.

## Results

### Restricted and constitutive expression of ZBP1 in normal and myeloma plasma cells

Although the role of ZBP1 in tumor biology is unknown, by searching for genes selectively expressed in MM we identified *ZBP1* as highly and selectively expressed in MM cell lines (MMCL) but not in other cancer cell lines **(Fig. 1a** and **Supplementary Fig. 1a)**. *ZBP1* is universally expressed in primary myeloma PC (n=767 patients; **Fig. 1b)** and at similar levels as the PC defining transcription factor *PRDM1* (BLIMP1). In human late mature B cells *ZBP1* expression was very low in germinal center B cells (GCB), detected in naive and memory B cells but PC displayed by far the highest expression **(Supplementary Fig. 1b).** We confirmed expression of *ZBP1* mRNA (qPCR) and/or protein (immunoblotting) in MMCL **(Fig. 1c, d)** and in primary myeloma PC **(Fig. 1e, f)**, but not in other hematopoietic or epithelial cancer cell lines **(Supplementary Fig. 1c, d)**, normal blood lineage cells **(Supplementary Fig. 1e, f)** or healthy non-hematopoietic tissues **(Supplementary Fig. 1g)**. Immunohistochemistry of human tonsils and lymph nodes showed expression of ZBP1 primarily in a group of PAX5^−^IRF4^+^ GCB cells i.e., those committed to PC differentiation^23,24^ and in interfollicular and subepithelial IRF4^+^ PC with low level expression in mantle zone B cells **(Fig. 1g and Supplementary Fig. 1h, i).** In the BM, expression of ZBP1 is mostly restricted in myeloma and normal PC but no other blood lineage cells **(Fig. 1h, and Supplementary Fig. 1h, j)**. These results show constitutive and restricted expression of ZBP1 in myeloma PC as well as in late GCB cells and normal PC.

**Fig. 1:**
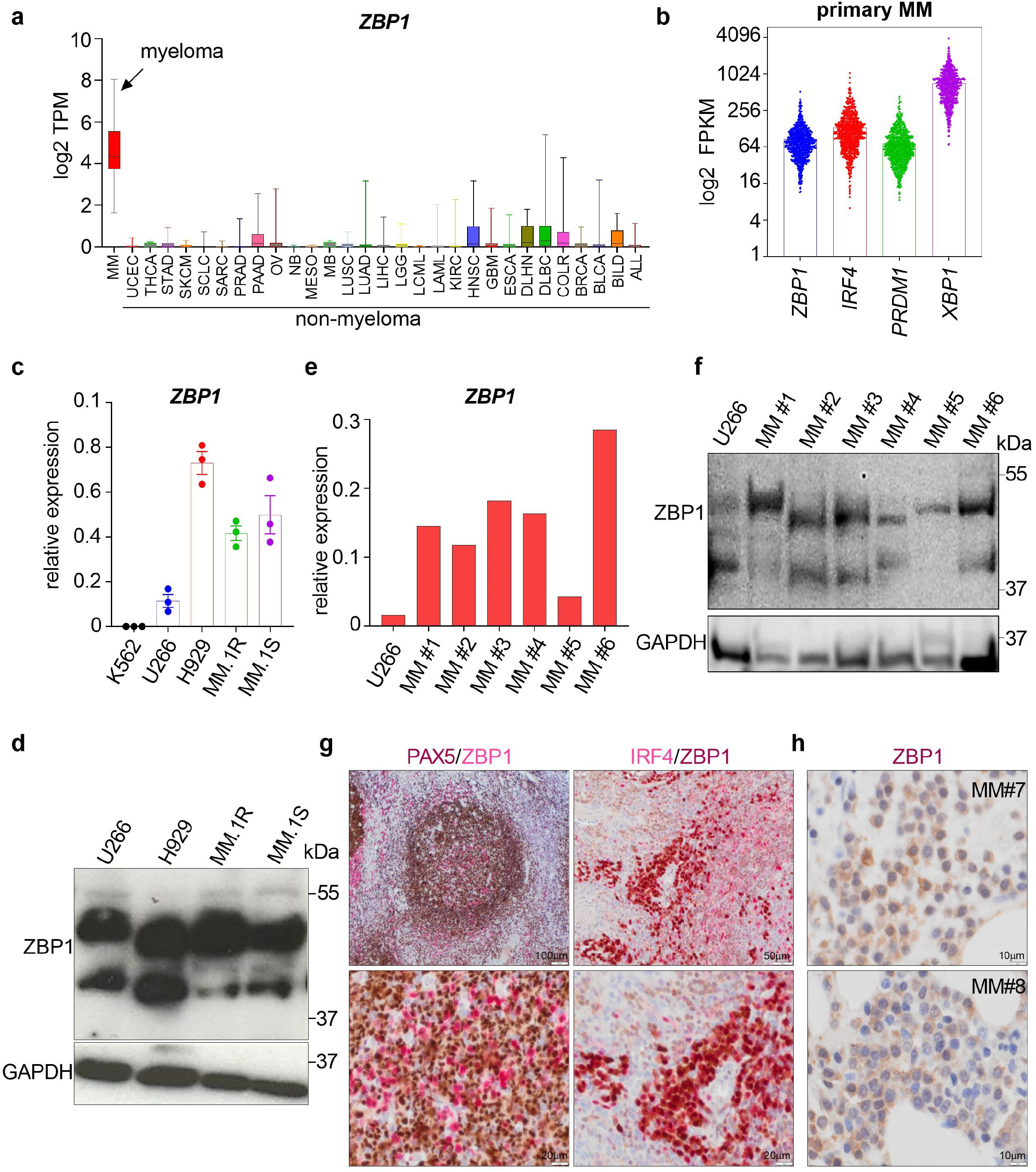
Restricted and constitutive ZBP1 expression in normal and myeloma plasma cells. **(a)** mRNA expression levels of *ZBP1* in >800 cancer cell lines including 27 MMCL. Box-whiskers plot shows minimum to maximum log2 TPM values. **(b)** mRNA expression levels of *ZBP1* in purified myeloma PC (n=767 MM patients) compared to PC-lineage defining transcription factors *IRF4*, *XBP1* and *PRDM1.* Bars show mean. **(c)** *ZBP1* mRNA expression as assessed by qPCR in 4 MMCL and in the erythromyeloid cell line K562 (n=3; data shown as mean ± standard error of mean-SEM). **(d)** ZBP1 expression in indicated MMCL as assessed by immunoblotting using GAPDH as loading control. Two main isoforms ~48 and ~40 kDa are detected in MMCL. **(e, f)** ZBP1 mRNA and protein expression in 6 purified myeloma bone marrow PC with U266 MMCL shown as control. **(g)** Immunohistochemistry on tonsil: Panel on the left depicts low and high magnification of a germinal center where expression of PAX5 and ZBP1 is mutually exclusive in most parts indicating ZBP1 expression in plasma cells that have downregulated PAX5. Panel on the right depicts sub-epithelial tonsillar plasma cells showing co-expression of IRF4 (MUM.1) and ZBP1. **(h)** ZBP1 expression in bone marrow myeloma PC from two patients with MM.

### ZBP1 is required for optimal T cell-dependent humoral immune responses in mice

Consistent with a conserved expression pattern, a previous transcriptome analysis of murine B cell development identified *Zbp1* as one of the signature genes that define transition from follicular B cells to plasmablasts and mature PC^25^ **(Fig. 2a).** To determine the role of ZBP1 in GCB and PC development and function, we assessed the humoral immune responses against the T cell-dependent antigen NP-KLH using alum as adjuvant^26^. We found that while *Zbp1* mRNA is undetectable in flow-sorted GCB cells and PC from *Zbp1*^−/−^ mice in response to either NP-KLH-alum or alum-only control, it increased from GCB cells to PC in wild type (WT) littermates and this increase was more pronounced upon immunization with NP-KLH-alum **(Fig. 2b)**.

**Fig. 2:**
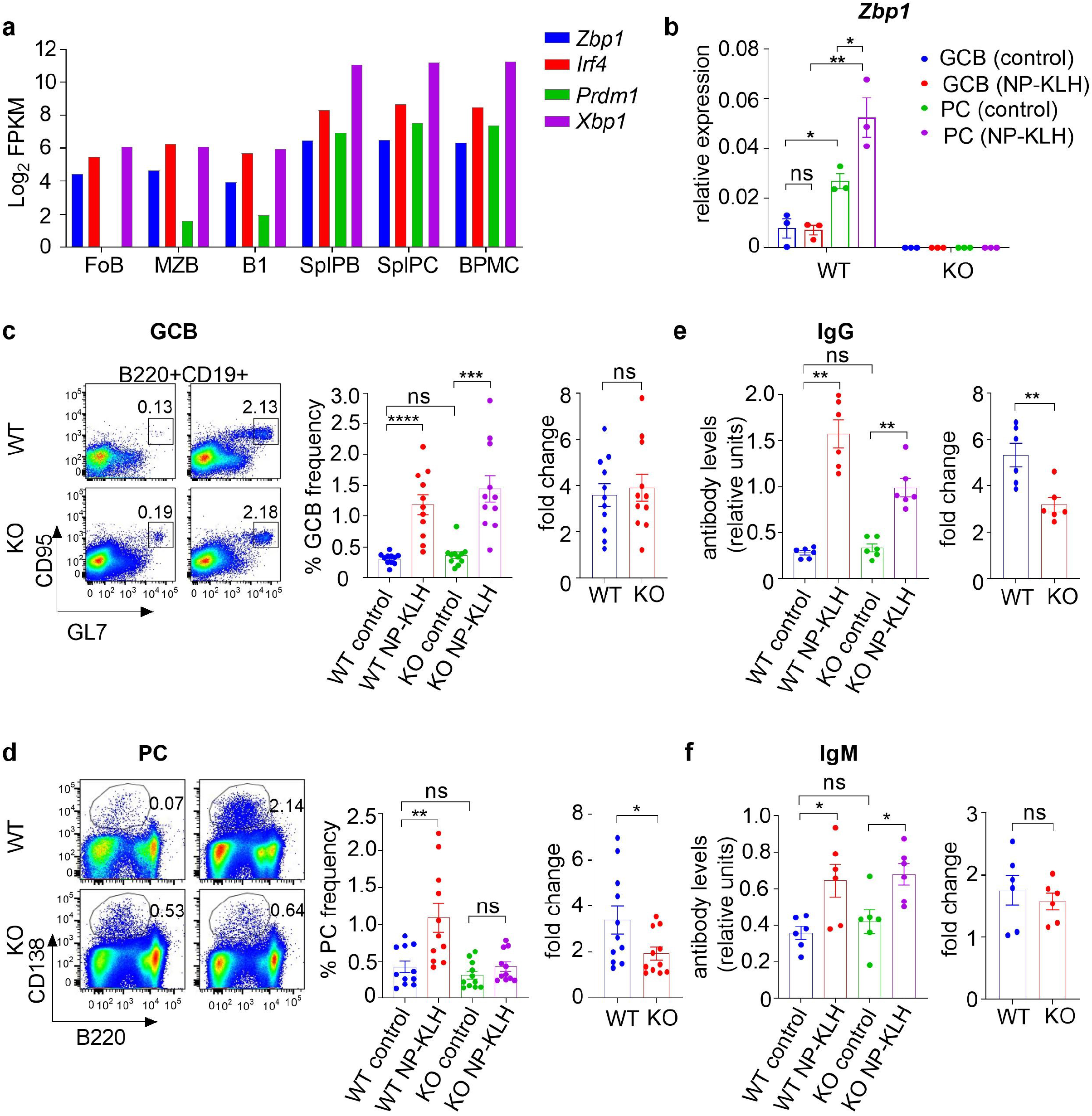
ZBP1 is required for optimal T cell-dependent humoral immune responses in mice. **(a)** mRNA expression levels of *Zbp1* in murine follicular B cells (FoB), marginal-zone B cells (MZB), splenic plasmablasts (splPB) and splenic and bone marrow plasma cells (splPC, BMPC) compared to PC-lineage defining transcription factors (figure created from previously published RNA-seq data^25^). **(b)** *Zbp1* mRNA levels as assessed by qPCR in flow-sorted splenic GCB and PC derived from *Zbp1*^−/−^ mice or their WT littermates immunised with just alum (control) or alum-NP-KLH (NP-KLH). (n=3). **(c)** Flow-cytometric identification of splenic GCB cells as B220^+^CD19^+^GL7^+^CD95^+^ and GCB frequency in *Zbp1*^−/−^ mice and their WT littermates immunised with just control or NP-KLH. Left bar graph: GCB cell frequency in *Zbp1*^−/−^ mice and their WT littermates in control-and NP-KLH-immunised animals. Right bar graph: GCB cell fold difference between WT and *Zbp1*^−/−^ mice after NP-KLH immunization normalised to median frequency of control animal. (n=11 mice/group). **(d)** Flow-cytometric identification of splenic PC as B220^lo^CD138^+^ and PC frequency in *Zbp1*^−/−^ mice and their WT littermates in control (left) and NP-KLH (right)-immunised animals. Left bar graph: PC frequency in *Zbp1*^−/−^ mice and their WT littermates in control and NP-KLH-immunised animals. Right bar graph: PC fold difference between WT and *Zbp1*^−/−^ mice after NP-KLH immunization normalised to median frequency of control animals (n=11 mice/group). **(e, f)** NP-KLH specific IgG and IgM responses in *Zbp1*^−/−^ mice and their WT littermates in control- and NP-KLH-immunised animals. Left panels: IgG and IgM Ab relative levels; Right panels: IgG and IgM fold difference after NP-KLH immunization normalised to median Ab levels of control animals (n=6 mice/group). Data shown as mean ± SEM, unpaired t-test; * p≤ 0.05, ** p≤ 0.01, *** p≤0.001, **** p≤ 0.0001, ns-not significant (p> 0.05).

Flow-cytometric analysis of splenic B220^+^CD19^+^GL7^+^CD95^+^ GCB cells and B220^lo^CD138^+^ PC frequency showed no baseline differences in *Zbp1*^−/−^ mice and their WT littermates and similarly GCB cell frequency was not significantly different in WT and *Zbp1*^−/−^ animals after NP-KLH-alum immunization **(Fig. 2c, d)**; however, the increase in PC frequency and antigen-specific IgG (but not IgM) serum levels in immunised *Zbp1*^−/−^ animals was significantly lower as compared to NP-KLH-alum immunised WT animals **(Fig. 2e, f)**. These findings suggest that although Zbp1 is not required for GCB cell and PC development under steady-state it is required for optimal, T cell-dependent humoral immune responses; whether cellular NA in complex with Zbp1 play a role in this process remains to be addressed.

### ZBP1 is required for myeloma cell proliferation and survival

To start exploring the functional role of ZBP1 in MM, we depleted its expression in MMCL by targeting both of its main isoforms, i.e., isoform 1 comprising Zα and Zβ domains and the shorter isoform 2 which lacks Zα. Depletion by shRNA-mediated knockdown **(Supplementary Fig. 2a-c)** of either ZBP1 isoform 1 (shRNA1) or both isoforms 1 and 2 (shRNA2) was toxic to all four MMCL tested **(Fig. 3a)** suggesting that the observed effect is mediated by depletion of isoform 1 and cannot be rescued by isoform 2. This effect was specific because we did not observe any significant effects on the growth of shRNA-transduced erythromyeloid K562 or epithelial HeLa cells **(Fig. 3b)** which lack expression of ZBP1 **(Supplementary Fig. 1c, 2d)**. Further, depletion of shRNA1-transduced myeloma cells was at least in part rescued by overexpression of *ZBP1* cDNA with appropriate silent mutations **(Supplementary Fig. 2e)** and it was not observed in myeloma cells transduced with a ‘seed’ (i.e., ‘off-target’) shRNA1 control^27^ that as expected did not impact on expression of ZBP1 **(Supplementary Fig. 2f, g).** Using a doxycycline-inducible shRNA system, we found that as well as *in vitro* **(Supplementary Fig. 2h, i)**, ZBP1 depletion also inhibited subcutaneous myeloma tumor growth *in vivo* **(Fig. 3c, d and Supplementary Fig. 2j, k)**. Together, these findings suggest an important role of ZBP1 and in particular of isoform 1 in myeloma cell proliferation.

**Fig. 3:**
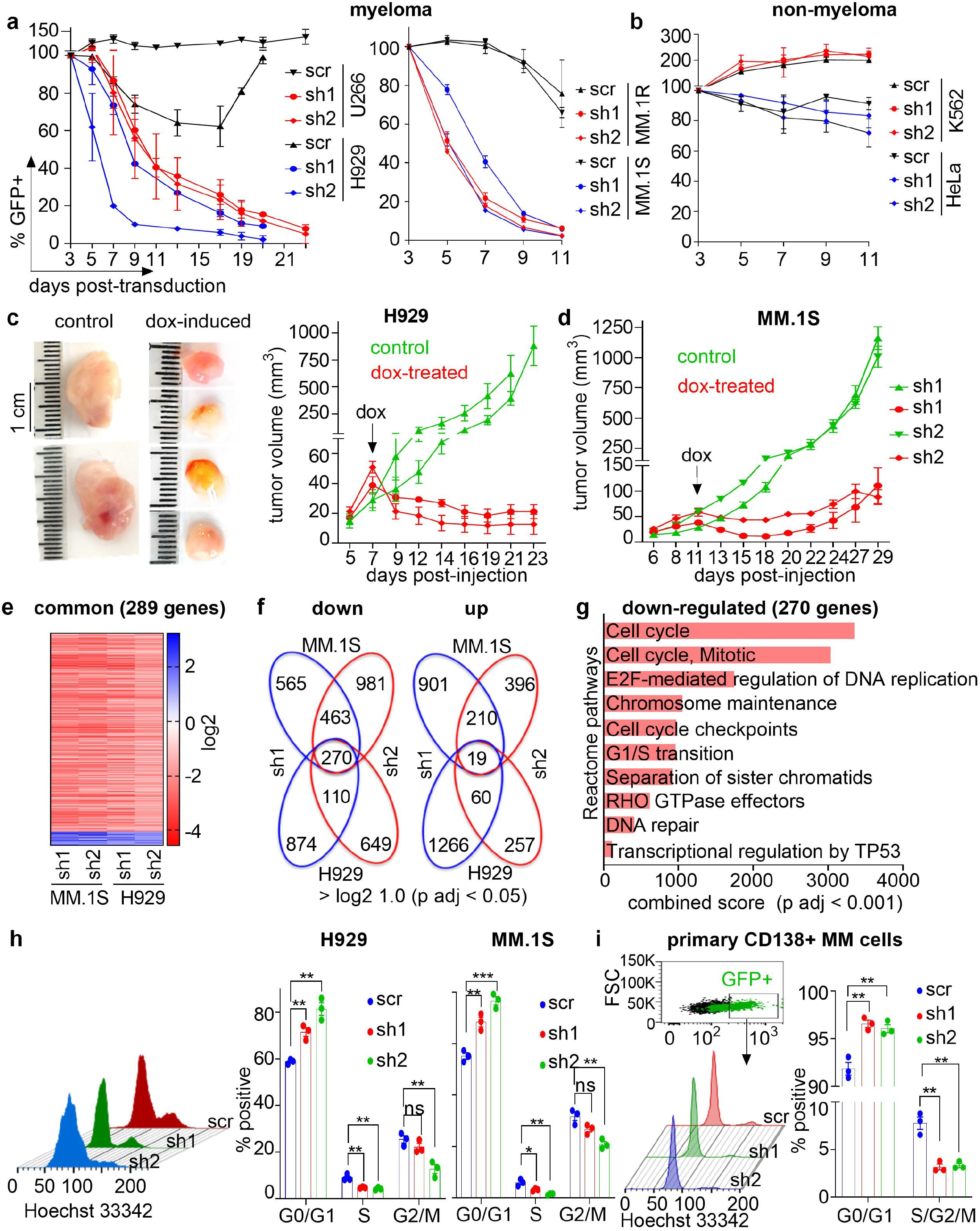
ZBP1 is required for myeloma cell proliferation and survival. **(a)** Percent GFP^+^ cells after transduction with GFP-encoding, *ZBP1*-targeting shRNA1 (sh1) or 2 (sh2) or appropriate scramble control (scr) lentiviral constructs. Data for 4 MMCL, normalised to day 3 GFP expression for each shRNA shown (n=3). **(b)** Percent GFP^+^ cells after transduction of K562 and HeLa cells, that do not express ZBP1, with ZBP1-targeting sh1 or sh2 or scr control (n=3). **(c, d)** Photographs of tumors explanted at sacrifice from control (i.e., non-Dox-treated) or Dox-treated animals engrafted with H929 myeloma cells transduced with Dox-inducible sh1 or sh2 targeting ZBP1 (c). Subcutaneous tumor volume of H929 or MM.1S over a period of up to 4 weeks in control or Dox-treated animals (c and d; n=4-5 mice/group). **(e)** Heatmap of the top 289 commonly and differentially expressed genes as assessed by RNA-seq shared by the two MMCL transduced with anti-ZBP1 sh1 or sh2 or scr control. **(f)** Venn diagram showing number of shared, differentially expressed genes in two MMCL treated with anti-ZBP1 shRNA1 or 2 or scr control. Replicate experiment, and commonly shared genes were identified based on log2 fold difference with cut off adjusted P-value (p adj) <0.05. **(g)** Enrichr pathway enrichment analysis of the shared 270 genes downregulated in two MMCL treated with anti-ZBP1 shRNA1 or shRNA2 or scr control. **(h)** Flow-cytometric analysis of cell cycle in MMCL H929 or MM.1S transduced with anti-ZBP1 shRNA1 or shRNA2 or scr control (left). Analysis was performed on day 4 post-transduction. Cumulative data from three independent experiments for each cell line, showing impact of ZBP1 depletion on cell cycle progression (right). **(i)** Flow-cytometric analysis of cell cycle in purified, bone marrow-derived myeloma PC (n=3 MM patients) transduced with anti-ZBP1 shRNA1 or shRNA2 or scr control. Data shown as mean ± SEM, unpaired t-test; ** p≤ 0.01, *** p≤0.001.

In line with this observation, transcriptome analysis of two ZBP1-depleted MMCL in which oncogenic transcriptomes are driven by the MAF (MM.1S) or MMSET (H929) oncogenes revealed that 270 genes that are significantly downregulated in both cell lines and by both shRNAs were enriched in cell cycle control pathways **(Fig. 3e-g, and Supplementary Fig. 3a)**.

In addition, we performed gene set enrichment analysis (GSEA) for each shRNA-related transcriptome in each cell line and in all four cases there was a consistent and significant enrichment for cell cycle related pathways amongst downregulated genes. We also identified significant enrichment for IFN type I pathway in upregulated genes induced by both shRNA1 and 2 in MM.1S cells and in downregulated genes by shRNA1 only in H929 cells **(Supplementary Fig. 3g).** This suggests that while ZBP1 mediates transcriptional repression of the IFN type I pathway in MM.1S cells, it retains its canonical function in H929. This disparate role of ZBP1 in the regulation of IFN type I response might reflect the distinct transcriptomes of MM.1S and H929 MMCL imposed by their primary driver oncogenes MAF and MMSET respectively.

Further, we validated reduction of the mRNA expression levels of the cell cycle regulators Ki-67, FOXM1 and E2F1 upon ZBP1 depletion by qPCR **(Supplementary Fig. 3a)** and also confirmed these findings at the protein level by immunoblotting (FOXM1 and E2F1) and flow-cytometry (Ki-67) **(Supplementary Fig. 3b-d)**. Indeed, supporting a critical role of ZBP1 in promoting myeloma cell proliferation and survival, flow-cytometric analysis of cell cycle status in ZBP1-depleted cells revealed arrest at G0/G1 phase **(Fig. 3h)** in conjunction with increased apoptosis as assessed by Annexin/DAPI staining **(Supplementary Fig. 3e)**. Notably, after lentivirally transducing purified patient-derived BM myeloma PC we confirmed that both *ZBP1* shRNA 1 and 2 induce cell cycle arrest **(Fig. 3i and Supplementary Fig. 3f),** thus confirming the role of ZBP1 in cell cycle regulation in primary myeloma PC as well as MMCL.

Previous work demonstrated that a transcriptional proliferative signature identifies a minority of MM patients with adverse prognosis^28,29^. Accordingly, GSEA of myeloma PC transcriptomes with top 5% highest vs 90% lowest *ZBP1* expression revealed significant enrichment in the former for cell cycle regulation pathways amongst over-expressed genes **(Supplementary Fig. 3h, i)**. Interestingly amongst these same over-expressed genes in ZBP1^hi^ patient subgroup, we also observed significant enrichment for IFN type I signalling suggesting that unlike in MM.1S cells, in primary myeloma PC, ZBP1 retains its canonical function in activating IFN type I response genes **(Supplementary Fig. 3h, i)**.

### ZBP1 as a scaffold for IRF3 constitutive activation by TBK1

We next investigated the downstream processes that might link constitutive ZBP1 expression in myeloma cells with regulation of cell cycle. Unlike in non-malignant cells in which phosphorylation and activation of IRF3 requires activation of sensors such as cGAS-STING and ZBP1 by cytosolic NA^9,11^, we found that IRF3 was constitutively phosphorylated (pIRF3) in myeloma cell lines **(Fig. 4a)**. In line with previous reports^30^, pIRF3 was also detected in other, non-myeloma cancer lines **(Fig. 4a)**. Importantly, as well as in MMCL, pIRF3 was also detected in primary BM myeloma CD138^+^ PC but not in non-malignant BM CD138^−^ cells from the same patients **(Fig. 4b)** thus establishing that IRF3 is constitutively phosphorylated in MM.

**Fig. 4:**
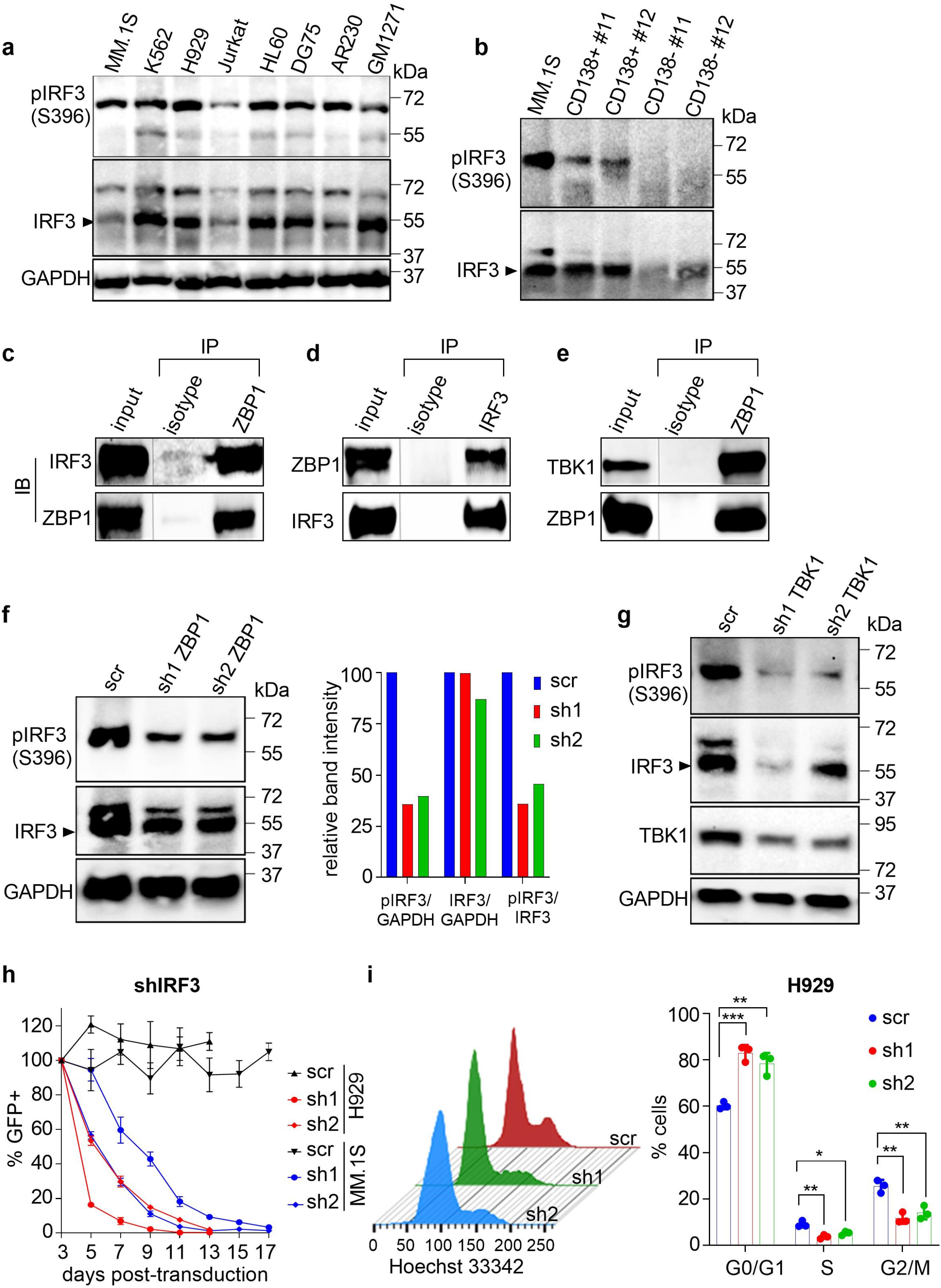
ZBP1 as a scaffold for IRF3 constitutive activation by TBK1. **(a)** Immunoblotting analysis of pIRF3/IRF3 expression in myeloma (MM.1S and H929) and non-myeloma cell lines (K562; Jurkat; HL60: acute myeloid leukemia; DG75: B lineage; AR230: chronic myeloid leukemia; GM1271: EBV-transformed B cell lineage). **(b)** pIRF3/IRF3 expression in purified myeloma bone marrow PC and non-myeloma bone marrow cells from two patients with MM. **(c, d)** Immunoblotting against IRF3 and ZBP1 in MM.1S cells following immunoprecipitation with anti-ZBP1 (c), -IRF3 (d) or isotype Ig Ab. **(e)** Immunoblotting against TBK1 and ZBP1 in MM.1S cells following immunoprecipitation with anti-ZBP1 or isotype Ig Ab. **(f)** pIRF3/IRF3 expression following anti-ZBP1 shRNA1 or shRNA2 or scr lentiviral transduction of MM.1S cells (left) and imageJ quantification of immunoblotting bands (right). **(g)** pIRF3/IRF3 expression following anti-TBK1 shRNA1 (sh1) or 2 (sh2) or scr transduction of MM.1S cells. **(h)** Percent GFP^+^ cells after transduction with *IRF3*-targeting shRNA1 or shRNA2 or scr control. Data for MMCL H929 and MM.1S cells, normalised to day 3 GFP expression for each shRNA shown (n=3). **(i)** Flow-cytometric analysis of cell cycle in MMCL transduced with anti-IRF3 shRNA1 or shRNA2 or scr control. Analysis was performed on day 4 post-transduction and cumulative data for H929 showing impact of IRF3 depletion on cell cycle progression in MMCL (n=3). Data shown as mean ± SEM, unpaired t-test; * p≤ 0.05, ** p≤ 0.01, *** p≤0.001.

ZBP1 interaction with IRF3 was previously only shown in an exogenous system^1,31^; using protein co-immunoprecipitation we found that endogenous ZBP1 interacts with IRF3 in MM.1S myeloma cells **(Fig. 4c, d)** and TBK1 **(Fig. 4e)**. Using co-transfection and different *ZBP1* deletion cDNA constructs **(Supplementary Fig. 4a)**, we found that ZBP1-IRF3 interaction requires the RHIM domain-containing C-terminus of ZBP1 and possibly the Zα and Zβ domains **(Supplementary Fig. 4b, c)**. Further, while in ZBP1-depleted cells total IRF3 and TBK1 levels were not appreciably altered, pIRF3 and pTBK1 levels markedly decreased thus showing a post-translational dependency of constitutive phosphorylation of IRF3 and TBK1 on ZBP1 **(Fig. 4f and Supplementary Fig. 4d)**. Finally, shRNA-mediated depletion of TBK1 resulted in decrease of IRF3 phosphorylation **(Fig. 4g)**. Together these data support a model whereby ZBP1 serves as a scaffold for TBK1-depedent constitutive phosphorylation of IRF3 in MMCL.

### IRF3 regulates cell cycle in myeloma cells

As observed for ZBP1, IRF3 depletion resulted in cell cycle arrest and apoptosis of myeloma cells **(Fig. 4h, i and Supplementary Fig. 5a-c)**, and depletion of TBK1 had a similar effect **(Supplementary Fig. 5d-g)**. In addition, transcriptome analysis of IRF3-depleted MM.1S myeloma cells revealed that amongst 185 genes downregulated by both anti-*IRF3* shRNA1 and shRNA2, 109 were also downregulated upon ZBP1 depletion **(Fig. 5a, b)** and these are also enriched for cell cycle regulation **(Fig. 5c).** Only 23 genes were shared amongst those upregulated upon depletion of both ZBP1 and IRF3.

**Fig. 5:**
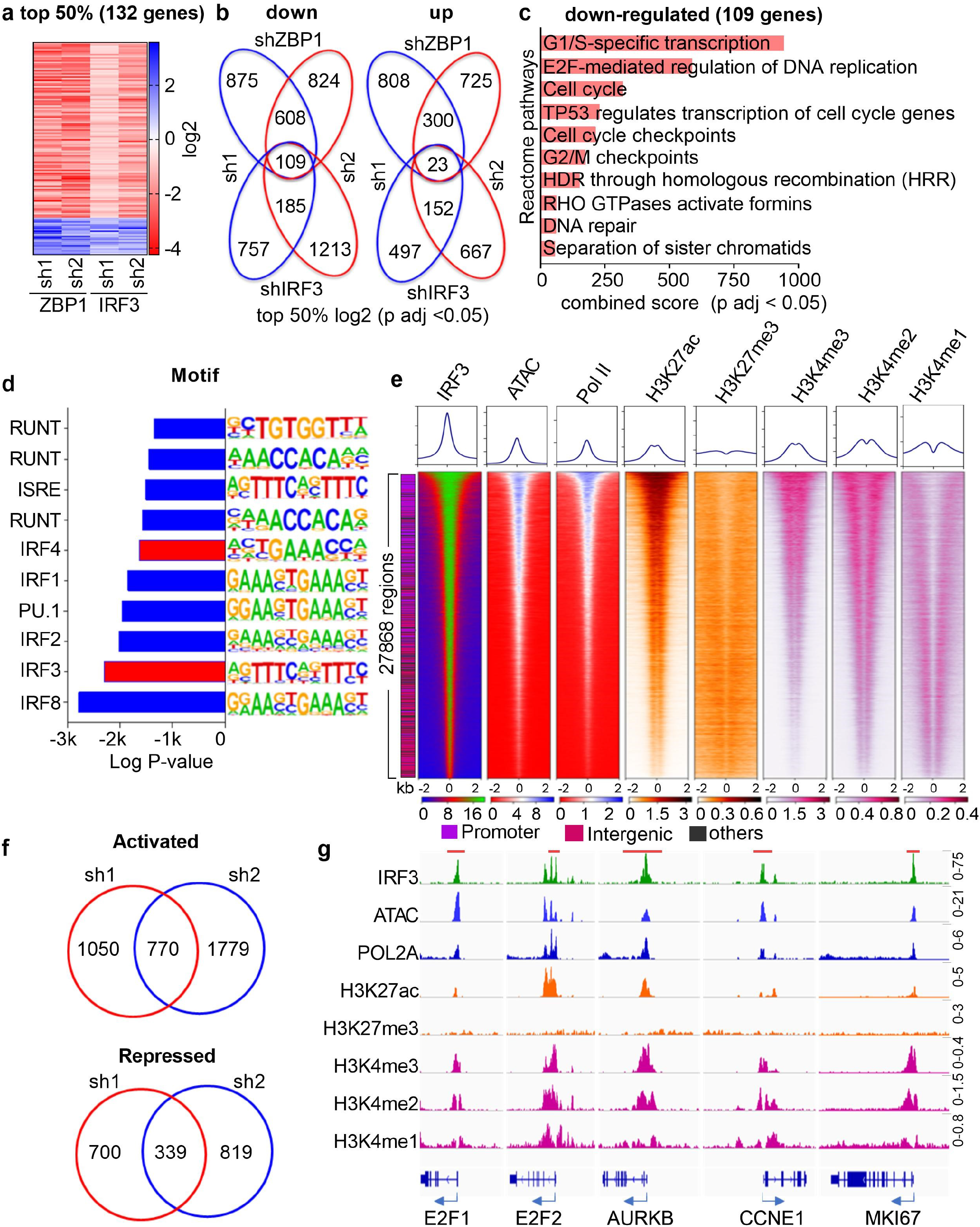
IRF3 regulates cell cycle in myeloma cells. **(a)** Heatmap showing differentially expressed genes upon intersection of top 50% differentially expressed genes following *ZBP1* or *IRF3* depletion in MM.1S myeloma cells. **(b)** Venn diagram showing number of common genes up- and -down regulated among top 50% of differentially regulated genes based on log_2_-fold change (p adj < 0.05) upon *ZBP1* or *IRF3* depletion in MM.1S myeloma cells. **(c)** Enrichr pathway enrichment analysis of the shared 109 genes downregulated in MM.1S cells upon *ZBP1* and *IRF3* depletion. **(d)** Transcription factor motif logos and enrichment at IRF3-bound genomic regions in MM.1S myeloma cells. **(e)** Metagene (top) and heatmap (bottom) representation of IRF3 genome-wide binding as assessed by ChIP-seq in MM.1S cells along with various histone marks, RNA Polymerase II(Pol II) binding and chromatin accessibility as assessed by ATAC-seq. Genomic feature annotation for each peak in the heatmap is shown on the left. IRF3 binding is observed in genomic regions marked for active transcription i.e., with increased chromatin accessibility, activating chromatin marks (H3K27ac, H3K4me1/3) and Pol II binding. **(f)** Venn diagram showing number of genes predicted to be directly regulated (activated or repressed) by IRF3 as assessed by integration of cistrome and post-*IRF3* depletion transcriptome using BETA-plus software. **(g)** IGV browser snapshots of IRF3 and Pol II binding, chromatin accessibility and histone mark enrichment at regulatory areas of several genes promoting cell cycle progression and cell proliferation. The red block on the top indicates 5kb.

To identify candidate transcriptional targets of IRF3 in myeloma cells we generated and mapped its genome-wide binding by ChIP-Seq in MM.1S cells. IRF3-bound regions (promoter, intergenic and intronic; **Supplementary Fig. 6a**) were highly enriched in IRF3 binding motifs **(Fig. 5d)**. In the same cells, we correlated chromatin accessibility, RNA Pol II binding, activating (H3K27ac and H3K4me1/2/3) and repressive (H3K27me3) histone marks **(Fig. 5e)** with genome-wide IRF3 binding. This showed that IRF3 binding occurs in nearly 28,000 highly accessible chromatin regions with activating transcriptional potential as revealed by Pol II binding and presence of activating histone marks. Thus, constitutively phosphorylated IRF3 in myeloma cells is highly transcriptionally active in the nucleus.

Next, to obtain the compendium of genes directly regulated by IRF3 in MM.1S myeloma cells, for each IRF3 shRNA we integrated the transcriptome of IRF3-depleted myeloma cells with the IRF3 cistrome using BETAplus software **(Fig. 5f and Supplementary Fig. 6d)**. After data intersection, we found that the 770 genes predicted to be directly activated by IRF3, included the key cell cycle regulators *E2F1*, *E2F2*, *AURKB*, *CCNE1* and *MKI67* and were significantly enriched in cell cycle regulation pathways; **(Fig. 5g and Supplementary Fig. 6b, e, f).** Notably, the 339 genes predicted to be repressed by IRF3 were not enriched for IFN type I response genes **(Fig. 5f and Supplementary Fig. 6c, g, h)**. Consistent with this, while IRF3 binds in the regulatory regions of *IFNA1* (but not of *IFNB1*) and *ISG15*, a hallmark type I IFN response gene, expression of these genes was not altered upon IRF3 depletion **(Supplementary Fig. 6g, h)**.

### IRF3 regulates *IRF4* and co-operates with IRF4 to regulate cell cycle in myeloma cells

IRF4 is a transcription factor critical for normal PC development^32^ while in myeloma PC it cooperates with MYC to establish a transcriptional circuitry to which myeloma cells are highly addicted to^33^. Since the IRF4 motif was amongst the top-most enriched in regions bound by IRF3 **(Fig. 5d),** we explored potential synergy between IRF3 and IRF4 by overlaying our in-house-generated genome-wide binding profile of IRF3 with that of previously published IRF4 ChIP-seq in MM.1S cells. First, we observed co-binding of the two transcription factors at the promoter and the previously established super-enhancer of IRF4^34^ **(Fig. 6a)**. This observation and the fact that *IRF4* expression is significantly downregulated following IRF3 as well as ZBP1 depletion **(Fig. 6b)** is consistent with IRF4 transcriptional regulation being downstream of ZBP1 and directly regulated by IRF3.

**Fig. 6:**
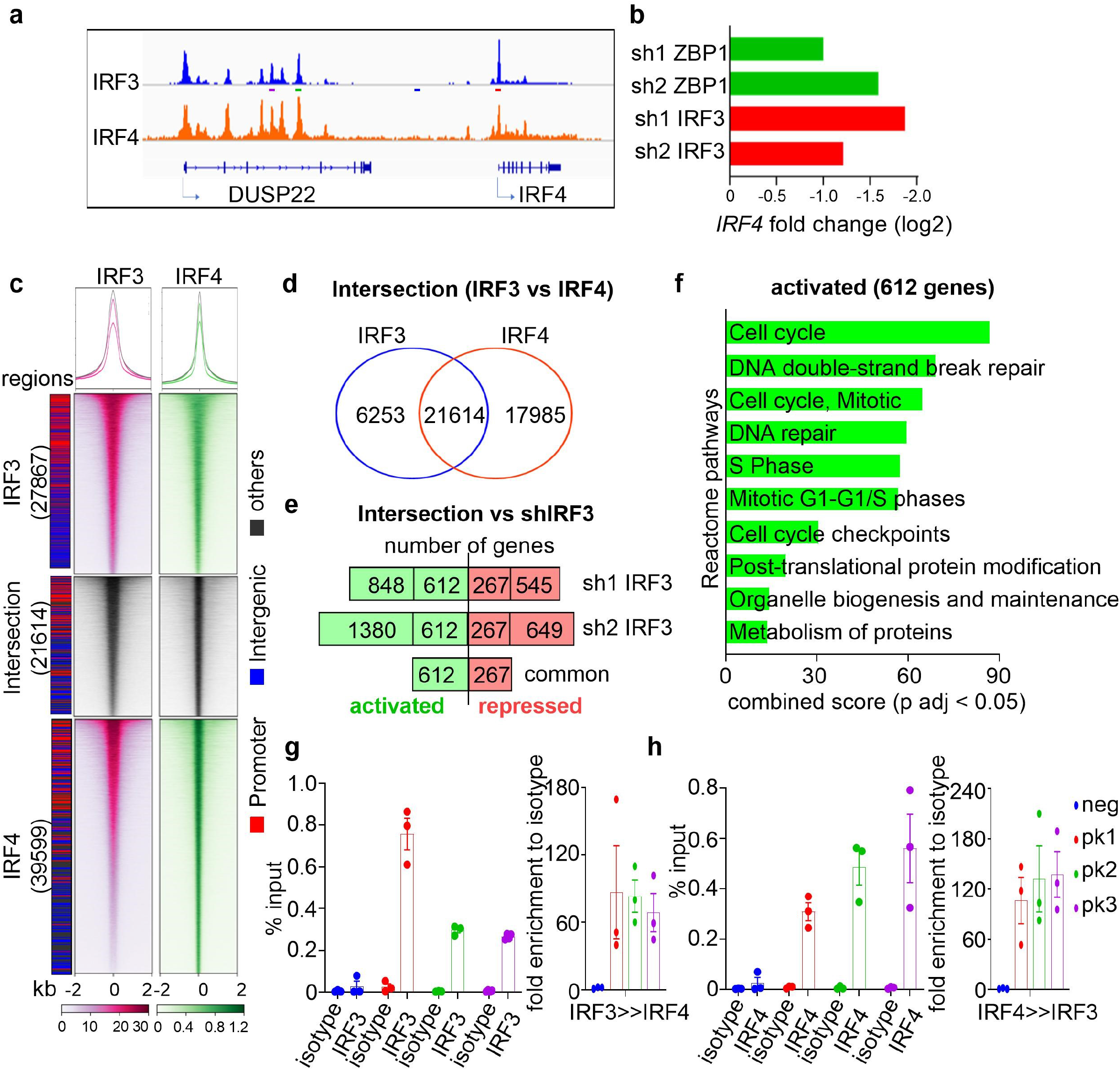
IRF3 co-operates with IRF4 and regulates cell cycle in myeloma cells. **(a, b)** IGV browser snapshots of IRF3 and IRF4 co-binding at the promoter and super-enhancer of IRF4 as assessed by ChIP-seq (a) and change in *IRF4* expression assessed by RNA-seq after depletion of indicated mRNA/protein by shRNA knock-down and in relation to scr control (b) in MM.1S cells (p adj <0.05). **(c)** Heatmaps of IRF3 and IRF4 binding and common binding regions of IRF3 and IRF4 (intersection). **(d)** Venn diagram showing number of genomic regions of IRF3 and IRF4 co-binding with gene regulatory potential as assessed by integration with whole transcriptome of *IRF3*-depleted MM.1S cells. For IRF3-IRF4 binding intersection the genome-wide binding of IRF3 and the top 50% IRF4 peaks with highest scores were included. **(e)** Number of genes predicted to be directly co-regulated (repressed or activated) by IRF3 and IRF4 binding. **(f)** Enrichr pathway enrichment analysis of genes predicted to be activated by IRF3-IRF4 co-binding. **(g)** Primary ChIP-qPCR against IRF3 (left) followed by re-ChIP-qPCR against IRF4 (right). Position of amplicons are shown as horizontal coloured bars in Fig. 6a. **(h)** Primary ChIP-qPCR against IRF4 (left) followed by re-ChIP-qPCR against IRF3 (right).

At a genome-wide level, 21,614 IRF3-bound chromatin regions were co-bound by IRF4 **(Fig. 6c, d)**. Correlating these with transcriptome changes following IRF3 depletion identified 612 and 267 genes predicted to be co-activated or co-repressed respectively by both IRF3 and IRF4 **(Fig. 6d, e)** with the former highly enriched in cell cycle regulators **(Fig. 6f)**. To validate this as an on-chromatin association of IRF3 with IRF4, we performed ChIP-re-ChIP assays at regions of IRF3-IRF4 co-binding using a region upstream of *IRF4* where no binding of either transcription factor was observed as a negative control. In all tested regions we found specific co-occupancy of IRF3 and IRF4 including at the *IRF4* promoter and super-enhancer and at genes regulating cell cycle, e.g., *E2F1*, *E2F2*, *MCM2* and *AURKB* **(Fig. 6g, h and Supplementary Fig. 7a, b)**.

## Discussion

Here we demonstrate that the NA sensor ZBP1 is an important and novel determinant of MM biology. We link the myeloma-selective constitutive expression of ZBP1 to constitutive IRF3 activation and regulation by pIRF3 of myeloma cell proliferation and IRF4 expression.

While other NA sensors such the DNA cGAS-STING and RNA RIG-1 or MDA-5 sensors are expressed constitutively and are activated upon NA binding^9,11,35^, ZBP1 differs in that as its expression is only detected in response to endogenous or exogenous NA, infection with viral pathogens or inflammatory stimuli including IFNs^1,6,3,36^. However, expression of ZBP1 has been studied only in a limited number of primarily murine cell lines and tissues. Our extensive analysis confirmed that ZBP1 expression is low or not detected in all human normal and cancer cells tested with the striking exception of cells in the late B cell development trajectory and in particular of plasma cells. Reflecting their cell of origin, we found high level constitutive ZBP1 expression also in MMCL and primary myeloma PC.

While our data does not address the molecular role of Zbp1 in late B cell development, at a cellular level it identifies an *in vivo* defect in *Zbp1*^−/−^ mice that consists of suboptimal humoral immunity in response to a well-established T cell-dependent antigen. The pattern of *Zbp1* expression, i.e., low in murine GCB but increasing in PC and more so upon immunisation suggests a conserved function that might be linked to the unique developmental features of PC such as production of large amounts of immunoglobulin mRNA upon transition from late GCB cells to mature PC.

We found that ZBP1 depletion had a profound and selective effect on MMCL proliferation and survival *in vitro* and *in vivo*. Similarly, in primary myeloma PC which are less proliferative than MMCL, depletion of ZBP1 also induced cell cycle arrest. Based on appropriate design of *ZBP1*-targeting shRNA we could determine that myeloma cell proliferation is sustained by the main isoform 1 that retains both Zα and Zβ domains but not isoform 2 that retains only domain Zβ. This is in line with *in vivo* studies showing that deletion of the Zα domain abrogates the physiological function of Zbp1^6,13,12^. Future research will explore the nature and origin of NA that are likely bound by the Zα and Zβ domains and their impact on the pro-proliferative function of ZBP1 in myeloma cells.

Transcriptomes of ZBP1-depleted myeloma cells driven by distinct primary oncogenes, i.e., MAF (MM.1S cells) and MMSET (H929 cells), highlighted cell cycle regulation as the prime pathway regulated by ZBP1. This novel pro-proliferative function of ZBP1 contrasts with the anti-proliferative potential of the IFN type I response, the canonical pathway regulated by ZBP1 in response to dsRNA and pathogens^3,6,37^. However, although GSEA suggested that in ZBP1-depleted MM.1S cells, ZBP1 mediates repression of the IFN type I response transcriptional programme, study of a large number of primary myeloma PC transcriptomes, as well as enrichment for cell cycle pathways amongst upregulated genes in ZBP1^hi^ myeloma PC, it also revealed a strong IFN type I response transcriptional signature. Indeed, tumor cell-intrinsic IFN type I responses have been documented in other cancers and have been shown to sensitise tumour cells to the loss of dsRNA editor ADAR^38^ while their suppression facilitates loss of immunosurveillance and breast cancer metastasis^39,40^. Together our findings in MM are consistent with a model whereby in the low proliferative, early disease myeloma PC, ZBP1 regulates in a balanced manner both anti- and pro-proliferative IFN type I and cell cycle gene pathways respectively. By contrast, in the highly proliferative MMCL, that are representative of advanced MM^41,42^ while the ZBP1-regulated IFN type I transcriptional programmes are attenuated or even repressed as in MM.1S cells, those supporting cell cycle progression persist and prevail. Consistent with this model, a recent comparison of primary myeloma PC and MMCL transcriptomes demonstrated enrichment for IFN type I response gene signature in myeloma PC from >700 patients at diagnosis while in relapsed disease myeloma PC and in MMCL, proliferative and not IFN type I gene signatures were dominant^42^.

One of the physiological roles of ZBP1 is to prevent or promote programmed cell death and inflammation through interaction with RIPK1 or RIPK3 respectively in response to pathogen or cellular dsRNA^4,5,7^. Alternatively, ZBP1 has been linked with activation of the IFN type I pathway via TBK1 phosphorylation of IRF3, with the latter directly activating amongst others the IFN genes themselves and *ISG15*, a hallmark IFN type I response gene^1,2,43^. However, such ZBP1-IRF3 interactions have only been shown in an exogenous system^1,31^. Here, for the first time we show interaction of endogenous ZBP1 with endogenous TBK1 and IRF3 in myeloma cells and phosphorylation of the latter by TBK1. ZBP1 depletion resulted in selective attenuation of IRF3 and TBK1 phosphorylation but no changes in the respective total proteins highlighting the role of ZBP1 as a physical platform that directs activation of TBK1 and IRF3.

While in response to inflammatory stimuli transcriptional activation by phosphorylation of IRF3 is expected to be transient, in myeloma cells, both primary and cell lines, we find that IRF3 is constitutively phosphorylated. This is not unique to MM since constitutively phosphorylated IRF3 has been reported in several ZBP1-negative cancer lines^30^ but not functionally investigated. By contrast, in myeloma cells we demonstrate that IRF3 binds to transcriptionally active regions of the genome and it directly regulates genes that promote cell cycle progression. Accordingly, IRF3 depletion in myeloma cells leads to cell cycle arrest and apoptosis. A similar pro-proliferative effect of IRF3 has been reported in acute myeloid leukaemia cells at a cellular level^44^. Of note, following integration of transcriptomes of IRF3-depleted MM.1S cells with the IRF3 cistrome, we found no evidence of IRF3 regulating IFN type I transcriptional programmes suggesting that ZBP1-associated repression of the IFN type I response in MM.1S cells is IRF3-independent. However, it is possible if not likely that in primary myeloma PC, as is the case for ZBP1, constitutively active IRF3 indeed activates IFN type I response genes along with proliferative pathways.

We also show that TBK1 depletion results in cell cycle arrest in myeloma cells. Although this cellular effect is likely linked to downstream regulation of cell cycle genes by pIRF3, since TBK1 is a pleiotropic kinase, other mechanisms are also possible. For example, TBK1 can regulate cell cycle by directly phosphorylating the centrosomal protein CP170, a critical component of the mitotic machinery^45^. TBK1 has been also shown to regulate cancer cell survival in co-operation with oncogenic K-RAS^19^. Interestingly, MM.1S and H929, the two MMCL in which TBK1 depletion results in cell cycle arrest, carry activating *K-RAS* and *N-RAS* mutations respectively (www.keatslab.org).

IRF4, the lineage-defining transcription factor in PC development, establishes an aberrant transcriptional circuity with MYC that renders myeloma PC highly dependent on a jointly regulated oncogenic programme that includes activation of cell cycle amongst other pathways^33^. Here, on the basis of IRF3 binding to the super-enhancer and promoter of IRF4 and the fact that depletion of IRF3 (and ZBP1) results in significant *IRF4* downregulation we demonstrate direct transcriptional activation of IRF4 by IRF3. Indeed, since as little as 50% reduction in *IRF4* expression levels is toxic to myeloma cells^33^, the >50% reduction in *IRF4* mRNA induced in IRF3-depleted cells would be expected to contribute significantly to myeloma cell death.

Our genome-wide and sequential ChIP assays demonstrate and validate extensive co-occupancy by IRF3 and IRF4 in the myeloma regulatory genome including at the super-enhancer of IRF4, with genes involved in cell cycle control being amongst the targets of the IRF3-IRF4 synergy. IRF3 binds chromatin either as a homodimer^46–48^ or in some cases as heterodimer in association with IRF7^49,50,48^. Similarly, IRF4 may function as a homodimer or in a heterodimeric association with PU.1 or IKZF1^51–53^; whether IRF3 and IRF4 heterodimerize with each other in myeloma cells remains to be investigated.

In summary, our data show that like other NA sensors, ZBP1 can regulate cellular pathways critical for cancer biology. Unlike the physiological transient activation of the ZBP1-IRF3 axis that comes about in response to dsRNA, ZBP1-IRF3 axis activation is constitutive and is co-opted into promoting proliferative pathways in myeloma PC and regulating expression of the critical myeloma oncogene IRF4. Guided by our initial delineation of the structural requirements of the ZBP1-IRF3 interaction in myeloma cells, disruption of the ZBP1-IRF3 axis will offer an opportunity for targeted and relatively selective therapeutic intervention in MM.

## Acknowledgments

We acknowledge funding from Blood Cancer UK (KP, NT, XX, VC) and KKLF (AK). We also acknowledge the Imperial NIHR Biomedical Research Centre, LMS/NIHR Imperial Biomedical Research Centre Flow Cytometry Facility, Imperial BRC Genomics Facility and the MRC/LMS Sequencing Facility for support.

## Author contributions

K.P and A.K conceived and designed the study. K.P and M.M.T performed CoIP and dox-inducible study *in vitro.* K.P and M.B performed shTBK1 study. P.T and K.N performed immunohistochemistry. A.C processed RNA-seq data for ZBP1. M.E.R processed RNA-seq and ChIP-seq data for ZBP1 and IRF3. K.P performed all other experiments *in vitro*, *in vivo* and integrated all the chip-seq and RNA-seq data and performed all other bioinformatics analysis and created all the figures. D.I provided erythroblast cells, V.S.C, A.K, N.T, X.X, I.V.K, I.R, H.W.A provided reagents. A.K supervised the study. K.P and A.K wrote the manuscript.

## Declaration of Interests

The authors declare no competing interest.

## Materials and methods

### *In vitro* cell lines and primary cell studies

Myeloma cell lines U266, NCI-H929 (DSMZ, Germany), MM.1R and MM.1S (ATCC, Manassas, VA, USA), were cultured in RPMI-1640 media (Sigma) supplemented with 10% FBS, 0.1mM non-essential amino acids, 1mM sodium pyruvate, 1% L-Glutamine and 1000U/ml Penicillin/Streptomycin at 5% CO_2_, 37°C. HeLa, HEK293T, DU145, MCF7 cells were cultured in DMEM (Sigma) and HCC95, SF295, LNCAP, K562, Jurkat and C1R cells in RPMI-1640 (Sigma) supplemented with 10% FBS, 1% L-Glutamine and 1000U/ml Penicillin/Streptomycin at 5% CO_2_, 37°C. Human bone marrow and peripheral blood cells from myeloma patients or healthy donor were collected under written consent and research ethical committee approval (REC reference number: 11/H0308/9). Plasma cells, T-cells, B-cells were isolated using CD138, CD3, CD19 antibody-coated magnetic beads (StemCell Technologies) respectively following manufacturer’s instructions. BM erythroblasts from normal donors were selected by flow-sorting (FACSAriall, BD Biosciences) after staining with anti-CD71 and anti-CD235a. Myeloma patient-derived CD138+ plasma cells were maintained in RPMI-1640 supplemented with 10ng/ml IL-6, 20% FBS, 0.1mM non-essential aminoacids, 1mM sodium pyruvate, 1% L-Glutamine and 1000U/ml Penicillin/Streptomycin at 5% CO_2_, 37°C.

### Cloning and lentiviral transduction

The puromycin selection gene was replaced with green fluorescent protein (eGFP) cDNA in Lentiviral pLKO.1 plasmid (Sigma). shRNA oligos (Sigma) were annealed by temperature ramp from 100°C to 25°C and cloned into pLKO.1 vector between AgeI and EcoRI sites. Doxycycline-inducible shRNAs were established using retroviral TRMPVIR vector (kind gift from Scott Lowe, Addgene plasmid #27994) as previously described^54^. ZBP1-full, ZBP1-ΔαΔβ cDNA constructs were kindly gifted by Stefan Rothenburg, University of California, USA^55^, and amplified ZBP1 cDNA using forward: GGGAATTCATGGCCCAGGCTCCTGCT and reverse: TAGCGGCCGCCTAAATCCCACCTCCCCA primers from pEGFPN.1 vector cloned into LeGO-iG2-IRES-EGFP vector (Addgene plasmid #27341) between EcoRI and NotI sites followed by 5x strep-tag II (TGGAGCCATCCGCAGTTTGAAAAA) sequences insertion between BamHI and EcoRI sites. ZBP1 ΔRHIM vector was constructed by amplifying strep-tagII-ZBP1 using forward: CCGGAATTAGGATCCATGTGGAGCCATCCG and reverse: TAGCGGCCGCCTAGGCTGACTTTGCTCTTC from LeGO-strep-tagII-ZBP1 full vector. The genetic rescue experiment was performed by co-expression of ZBP1 cDNA with mutation at shRNA1 binding regions 5’-CAAGAGGGAGCTCAA**TC**A**G**GT**AT**T**A**TA**TA**GAATGAA**G**AAGGAGTTGAAAGTCTCCCT-3’ (sh1 ZBP1 target site and **mutated sequences**) using LeGO-iC2 plasmid (addgene #27345) and sh1 ZBP1 in MM.1S cells. All viral particles were produced by calcium phosphate co-transfection of pRSV.REV, pMDLgpRRE and pMD2.VSVG (lentiviral) plasmids in HEK293T cells and concentrated by ultracentrifugation at 23,000 rpm for 100 minutes at 4°C. Myeloma and non-myeloma cells were treated with 8μg/ml polybrene (Sigma) and transduced with lentivirus by spinoculation at 2000 rpm, 37°C for 1hr followed by replacement of polybrene-media with appropriate culture media 24hr post-spinoculation. Details of shRNA sequences are listed in table 1.

TRMPVIR-sh1 ZBP1 (5’-3’): TCGAGAAGGTATATTGCTGTTGACAGTGAGCGCCAAGTCCTCTACCGAATGAAATAGTGAAGCCACAGATGTATTTCATTCGGTAGAGGACTTGGTGCCTACTGCCTCGG

TRMPVIR-sh2 ZBP1 (5’-3’): TCGAGAAGGTATATTGCTGTTGACAGTGAGCGGCACAATCCAATCAACATGATTAGTGAAGCCACAGATGTAATCATGTTGATTGGATTGTGCTGCCTACTGCCTCGG

**Table 1.**
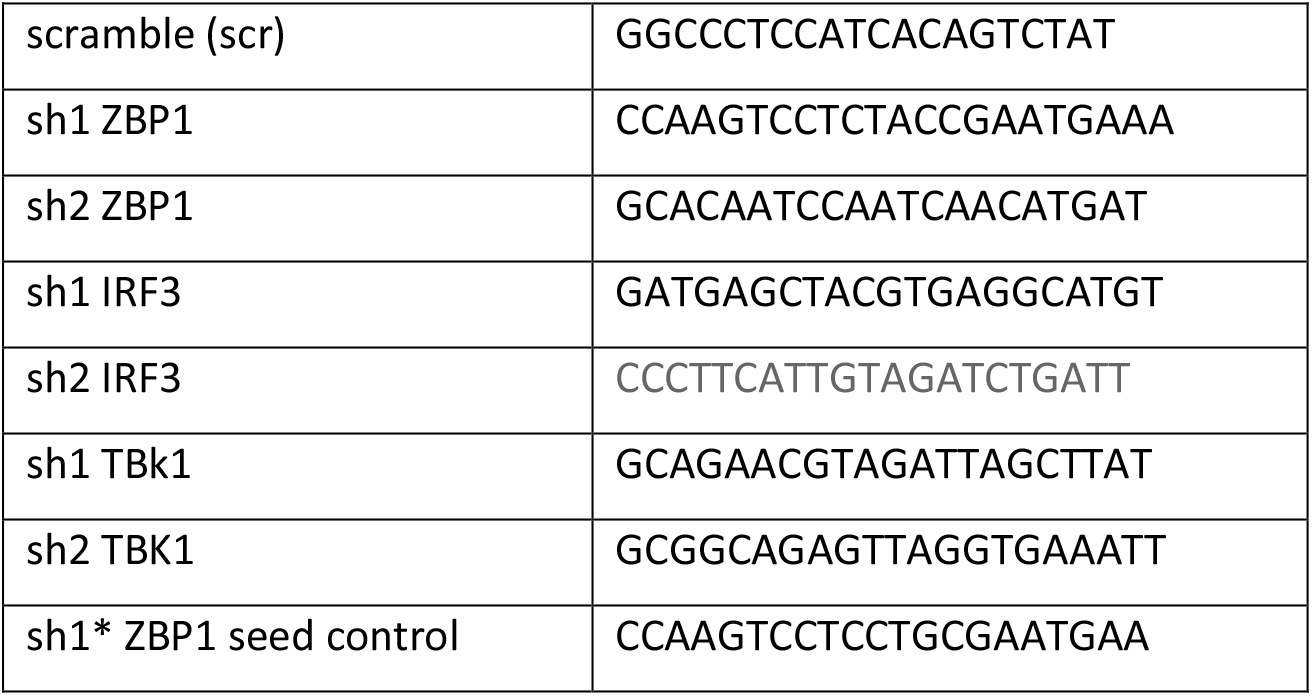
shRNA sequences (5’-3’)

### Cell cycle, cell proliferation and apoptosis assays

Cells were incubated with 10μM Hoechst 33342 stain in RPMI-1640 media supplemented with 2% FBS at 5% CO_2_, 37°C for 1 hour and cell cycle status of live cells was analysed by flow cytometry. Fluorescent marker eGFP frequencies of transduced cells were measured post-transduction by flow cytometry and cell proliferation was calculated based on the eGFP frequency on each day normalized to day 3 post-transduction applying the formula (percentage of live cells/percentage of total cells*eGFP frequency of live cells in the flow cytometry plot). For doxycycline-inducible shRNA cells, dsRed+ GFP+ (double positive) or control GFP+ only frequency was measured and followed the above procedure. Similarly, GFP+ mCherry+ (double positive) frequency was measured for genetic rescue experiment. Cells were stained with annexin V (Biolegend) according to manufacturer’s instructions followed by assessment of cell death by flow cytometry.

### Subcutaneous myeloma mouse model and immunization studies *in vivo*

Myeloma cells transduced with doxycycline-inducible shRNA-targeting *ZBP1* were resuspended in 150μl Matrigel Basement Membrane Matrix, LDEV-Free (Corning) and 10 million cells injected into NOD/SCIDgamma (NSG) mice subcutaneously. Tumor size was measured using a Vernier caliper and applied the formula (1/2(length × width^2^)) to obtain tumor volume. Upon a measurable tumor growth, 100μg/ml doxycycline added to drinking water and 0.2mg/kg doxycycline injected intraperitoneally and sacrificed all mice when tumor grown up to 15mm in any direction of the control mice. All the animals were kept in individually ventilated cages and studies were done under the Home Office project licence number PPL70/8586.

*Zbp1*^−/−^ animals^56^, already cross-bred to C57BL/6 animals for 4-5 generations were obtained from Manolis Pasparakis, Institute of Genetics, Cologne, Germany. They were further cross-bred with wild type C57BL/6 mice for another three generations and their littermates used to study T-cell dependent humoral immune response to 4-Hydroxy-3-nitrophenylacetyl hapten conjugated to Keyhole Limpet Hemocyanin (NP-KLH) antigen (Santacruz Biotech). 6mg/kg NP-KLH prepared in Imject™ Alum Adjuvant (Thermoscientific) 3:1 ratio and injected intraperitoneally into 10-12 weaks old age-matched *Zbp1*^−/−^ and wild type littermates. On day 4 post-immunization, 4mg/kg NP-KLH alone injected as booster dose and after 10 days post-immunization, blood samples were collected, and spleen was harvested. Single cell suspension of spleen cells were stained for B220 (BioLegend; clone: RA3-6B2), CD19 (BioLegend; clone: 6D5), CD95 (eBioscience; clone number: 15A7), GL7 (BioLegend; clone;GL), CD138 (BioLegend; clone: 281-2) and analyzed for germinal center activated B cells (GCB), plasma cell (PC) development. GCB (B220^+^ CD19^+^ GL7^+^ CD95^+^) and PC (B220^lo^CD138^+^) spleen cells were sorted using (FACS Aria) and total RNA was isolated and quantified *Zbp1* mRNA levels as described below. A standard ELISA method was used to quantify NP-KLH-specific IgG or IgM antibodies. Diluted (1:1000) serum samples used to detect levels of IgG by anti-IgG-HRP antibody (Bio-Techne) or IgM by anti-IgM-HRP antibody (Sigma) on 100μg/ml NP-KLH-coated ^57^. The antibody levels of immunized mice sera were normalized to their appropriate control alum-only immunized mice sera.

### qPCR

Cells were lysed and cellular RNA was isolated using Nucleospin RNA kit (Macherey-Nagel) followed by cDNA synthesis of 0.5-1μg total cell RNA using RevertAid cDNA synthesis kit (Thermoscientific). All specific cDNAs were quantified using Taqman probes (Applied Biosystems) or SYBR Select Master Mix in AB StepOne Plus Real-Time PCR (Applied Biosystems). *GAPDH* was used as a control to normalize the gene expression levels. Human ZBP1 qPCR (5’-3’): Forward: GCCAACAACGGGAGGAAGA; Reverse: ATCTTCTGGGCGGTAAATCGT

### RNA sequencing

Total RNA was isolated from eGFP-sorted myeloma cells transduced with either scramble or shRNA, targeting *ZBP1* or *IRF3* on day 4 post-transduction when >80% *ZBP1* or *IRF3* knock-down achieved, using Nucleospin RNA kit (Macherey-Nagel). The RNA quality was assessed in Bioanalyzer using the RNA pico kit (Agilent). 200ng of total RNA with RIN >9.5 were used to prepare library using NEBNext poly(A) mRNA Magnetic Isolation Module and the NEB Next Ultra II RNA library prep kits for Illumina (New Engand Biolabs). Qubit High Sensitivity DNA kit (Life Technologies) was used to quantify the libraries and Bioanalyzer High Sensitivity DNA kit (Agilent) for quality and fragment size assessment. 2nM of 350-400bp DNA library was sequenced using Illumina HiSeq 2500 (for *ZBP1*-depleted transcriptome) or NextSeq500 (for *IRF3*-depleted transcriptome) platform to obtain paired-end 150bp reads.

### ChIP-seq and ChIP-re-ChIP

MM.1S cells were cross-linked with 1% formaldehyde (Alfa Aesar) at 10^6^ cells/ml density for 15min at room temperature with gentle mixing followed by addition of 0.125M Glycine to final volume for 5min at room temperature with gentle mixing. Cells were washed thrice with ice cold 1x PBS with 10mM phenylmethylsulfonyl fluoride (PMSF) and 10^8^ cells were lysed with hypotonic lysis buffer (10 mM Hepes-KOH, pH 7.8, 10 mM KCl, 0.1 mM EDTA, and 0.1% IGEPAL CA-630) for 15minutes on ice followed by centrifugation at 5000g for 5min. Further the cell pellet was lysed in nuclear lysis buffer (1% SDS, 50mM Tris-HCl pH 8.0, 10mM EDTA pH 8.0, 300mM NaCl supplemented with 1x halt protease & phosphatase Inhibitor cocktail (Fisher scientific) for 15min on ice. The lysate was diluted 10 times with dilution buffer (0.01% SDS, 1 % Triton X-100, 1mM EDTA, 50mM Tris-HCl pH 8.0, 150mM NaCl) and sonicated to shear the chromatin DNA up to 500bp size. The lysates were precleared with 50 μl protein A/G magnetic beads (Life Technologies) and then IRF3-bound chromatins were pulled overnight, rotating at 4°C with either μ5g IRF3 antibody (BioLegend, clone:12A4A35) or equivalent isotype control conjugated with protein A/G magnetic beads. Immunoprecipitated beads were washed twice with wash buffer A (0.1% SDS, 1% TritonX-100, 1mM EDTA, 10mM Trish-HCl pH 8.0, 150mM NaCl), buffer B (0.1% SDS, 1% TritonX-100, 1mM EDTA, 10mM Trish-HCl pH 8.0, 500mM NaCl) and buffer C (0.25M LiCl, 1% IGEPAL CA-630, 1% sodium deoxycholate, 1mM EDTA, 10mM Tris-HCl pH 8.0) for 5min, rotating at 4°C. The ChIP complex was treated with 10mg/ml RNase A, 20mg/ml proteinase K and reverse crosslinked with a buffer containing 1% SDS, 50mM Tris HCl pH 8.0, 4M NaCl, 1mM EDTA at 65°C overnight. ChIP DNA was collected with Ampure XP beads (Beckman) and quantified using Qubit High Sensitivity DNA kit (Life Technologies). 1ng of ChIP DNA was taken to prepare library using NEBNext kit for Illumina (New Engand Biolabs) following manufacturer’s instructions and the quality or fragment size was assessed using the Bioanalyser High Sensitivity DNA kit (Agilent). 2nM of 400-500bp DNA library was sequenced using Illumina NextSeq500 platform to obtain paired-end 150bp reads.

For ChIP-reChIP, above protocol to pull IRF3 or IRF4-bound chromatin using IRF3 antibody (BioLegend, clone:12A4A35) or IRF4 antibody (BioLegend, clone: IRF4.3E4) and their equivalent isotype control respectively was followed. The chromatin was eluted in 1% SDS with 1x halt protease & phosphatase Inhibitor cocktail (Fisher Scientific) followed by 10 times dilution with elution buffer and repeated ChIP with the appropriate antibody. The ChIP-reChiP DNA was quantified using Qubit High Sensitivity DNA kit (Life Technologies) and quantified specific DNA fragments by qPCR, the primers sequences are listed in table 2.

**Table 2.**
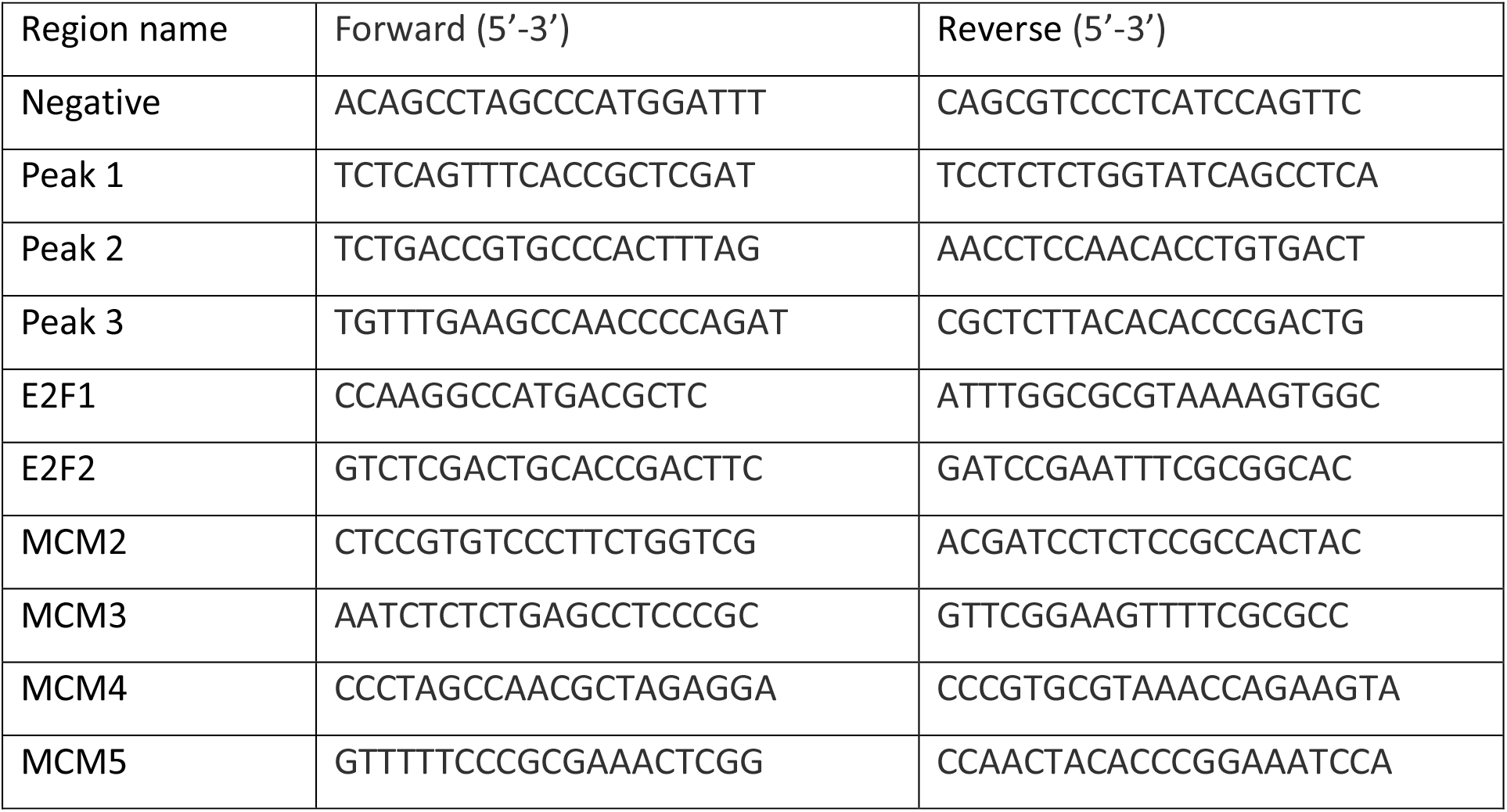
ChIP qPCR primers.

### Co-immunoprecipitation and immunoblotting

Transfected HEK293T cells were washed with ice-cold 1x PBS and 1×10^6^ cells were immediately lysed with co-immunoprecipitation (co-IP) buffer (25mM Tris buffer pH 7.4, 16mM NaCl, 0.5% Triton-X, 10% Glycerol, 2.5mM MgCl_2_, 0.1mM CaCl_2_, 1mM DTT) with 1x halt protease & phosphatase Inhibitor cocktail (Fisher scientific) for 30min on ice followed by centrifugation at 13000 rpm for 15min. The supernatant was incubated with 5% anti-strep-tag II-magnetic bead slurry (IBA) for overnight at 4°C with gently rotating. Beads were washed twice with co-IP wash buffer (25mM Hepes pH 7.4, 150mM NaCl, 1x halt protease & phosphatase Inhibitor cocktail (Fisher scientific) for 5min, 4°C. The IP-beads were collected under magnetic field and proceeded to immunoblot. For endogenous protein co-IP, followed the above protocol but immunoprecipitated with 2μg anti-ZBP1 (ThermoFisher Scientific; catalog number: PA5-20455), 2μg anti-V5-Tag (ThermoFisher Scientific; catalog number: 37-7500), 2μg anti-IRF3 (BioLegend; clone number: 12A4A35) and their equivalent isotype control antibodies conjugated with protein A/G magnetic beads.

For immunoblot, total cell protein was extracted by lysing the cells with buffer containing 250mM NaCl, 1.5mM MgCl_2_, 20mM HEPES pH 7.4, 0.5mM EDTA, 1% IGEPAL CA-630, 1% Triton X-100, 0.1% SDS, 10mM of PMSF (Sigma) supplemented with 1x halt protease & phosphatase Inhibitor cocktail (Fisher scientific) for 30min at 4°C followed by centrifugation at 13000 rpm for 10min at 4°C. samples were denatured in 1x LDS Sample Buffer (ThermoFisher Scientific) at 100°C for 7minutes. Total protein was quantified using BCA protein assay kit (ThermoFisher Scientific) according to manufacturer’s instructions and about 20μg of denatured total cell protein resolved in 10% SDS-PAGE and transferred to PVDF membrane followed by blocking with 5% BSA and antibody probing in 2.5% BSA solution with the indicated antibodies, anti-ZBP1 (1:1000; SantCruz Biotech; sc-67259) for Fig. 1d and Supplementary Fig. 1c, 1d, 1f and 2b however this product was discontinued meanwhile. Therefore, anti-ZBP1 (1:1000; ThermoFisher Scientific; catalog number: PA5-20455) antibody was used for further experiments. Of note, both anti-ZBP1 antibodies detected two main ZBP1 isoforms in myeloma cells. The anti-IRF3 (1:500; BioLegend; clone number: 12A4A35), anti-TBK1 (1:1000; SantCruz Biotech; sc-52957), anti-pTBK1(ser172) (1:250; Cell Signalling Technolog; catalog number: 5483), anti-V5-tag (1:500; Cell Signalling Technology; catalog number:13202S), anti-pIRF3(S396) (1:500; ThermoFisher Scientific; Catalog number: 720012), GAPDH (1:2000; SantCruz Biotech; sc-47724).

### Immunohistochemistry

Immunohistochemistry was undertaken on one-micron formalin-fixed paraffin-embedded tissue sections on Leica Bond III automated immunohistochemistry stainer. Prior to the procedure, paraffin sections were placed in oven preheated to 60°C for 30 minutes. Ready to use antibodies were used for PAX5 (clone number: PA0552; 15 minutes incubation) and MUM.1 (IRF4) (clone number: PA0129; 15 minutes incubation) antigens. Antibodies for ZBP1 (Sigma-Aldrich; catalogue number: HPA041256; 20 minutes incubation), CD3e (clone: NCC-L-CD3-565; 20 minutes incubation) and CD21 (clone: NCC-L-CD21-269; 20 minutes incubation) were diluted 1:100. 1:100 and 1:25 respectively prior to incubation. Heat induced epitope retrieval was undertaken for 20 minutes (PAX5 (EDTA buffer), MUM.1 (EDTA buffer), ZBP1 (citrate buffer) and CD21 (citrate buffer)) and 30 minutes (CD3e (citrate buffer)).

Signal detection for single immunostains was performed using Bond Polymer Detection Kit (clone number: DS9800) with DAB (brown colour) as the chromogen. For double immunostains, the sections were initially stained for CD3e, CD21, MUM.1 or PAX5 antigens as for single immunostaining protocol. This was followed by a sequential step for staining for ZBP1 antigen, where is the ZBP1 signals were detected using Bond Polymer Refine Red Detection Kit (Red signals; catalogue number: DS9390).

### Bioinformatics

For RNA sequencing analysis, reads were aligned and transcripts were quantified using STAR (v2.5.3a)^58^, for shRNAs targeting *ZBP1* in MM.1S and H929 cells; related to figure 2, against GRCh38 release 79 or with Salmon (v0.12.0)^59^, for shRNAs targeting *ZBP1* or *IRF3* in MM.1S; related to figure 4, against GRCh38 Gencode v28 transcript annotations. Differential expression analysis was performed in R (R Core Team, 2020 (https://www.R-project.org/)) with DESeq2 (v1.24.0) from STAR output or Salmon output using tximport (v1.12.3), and limma-voom for processing CCLE data with cut off p adj <0.05^58,60,61^. IRF3 ChIP sequencing reads were aligned with BWA MEM (v0.7.15) to GRCh38 genome (Gencode v28) with standard settings^62^. QC and duplicate marking were performed with Picard (v2.6.0) and samtools (v1.2). Tracks were generated with Deeptools (v3.3.1)^63^, and peaks were called with MACS2 (v2.1.1)^64^. Motif enrichment was performed with Homer (v4.10)^65^. The peaks were visualized using Integrative Genomes Browser (IGV) (v2.5.2)^66^. Binding and Expression Target Analysis (BETA)-plus package (v1.0.7)^67^ was used to integrate IRF3 cistrome, with the peaks within 2kb distance to TSS, and a complete transcriptome of *IRF3*-depleted MM.1S cells with cut off padj <0.05. Bigwig and BED files of ATAC-seq and Pol II, H3K27ac, H3K27me3, H3K4me3, H3K4me2, H3K4me1 and IRF4 ChIP-seq files of MM.1S cells were collected from Cistrome Data Browser^68^.

Bedtools (v2.25.0) Intersect^69^ was used to identify the common genome-wide binding of IRF3 and IRF4 factors. Deeptools computeMatrix and plotHeatmap (v3.4.1)^63^ with 2kb distance in reference to center of the region were used to visualize genome-wide binding of histone marks, Pol II, with IRF3, and IRF4 transcription factors. Homer (v4.10)^65^ was used for annotation of the genomic regions that are plotted in the heatmap. Gene Set Enrichment Analysis (GSEA) (v4.0.3) software^70^ was used for pathway annotation of differentially regulated genes with p adj <0.05 to analyse the pathways of Hallmark gene sets. Enrichr online web tool^71^ was used for pathways enrichment analysis of differentially regulated genes from RNA-seq and output of BETA-plus for integration of ChIP-seq and RNA-seq data. Significant pathways enriched with p adj < 0.05 were selected to create the figures and listed in the tables.

### Statistical analysis

Data graph and statistical analysis were performed using GraphPad Prism 8.0 software under institute licence. All experiments were repeated at least three times except for RNA-seq and ChIP-seq which were performed in replicates. Fold changes for *in vivo* data that were obtained in different time points were calculated by comparing the immunized groups to median value of control group. Comparison of two groups were performed using unpaired Student t-test. All the information on sample size, replicates, statistical method and significance are indicated in the figure legends. GraphPad Prism 8.0 or Morpheus (https://software.broadinstitute.org/morpheus) was used for heatmap creation with log2 transformed values of RNA-seq data.

---

**Supplementary Fig. 1 (related to Fig. 1):**
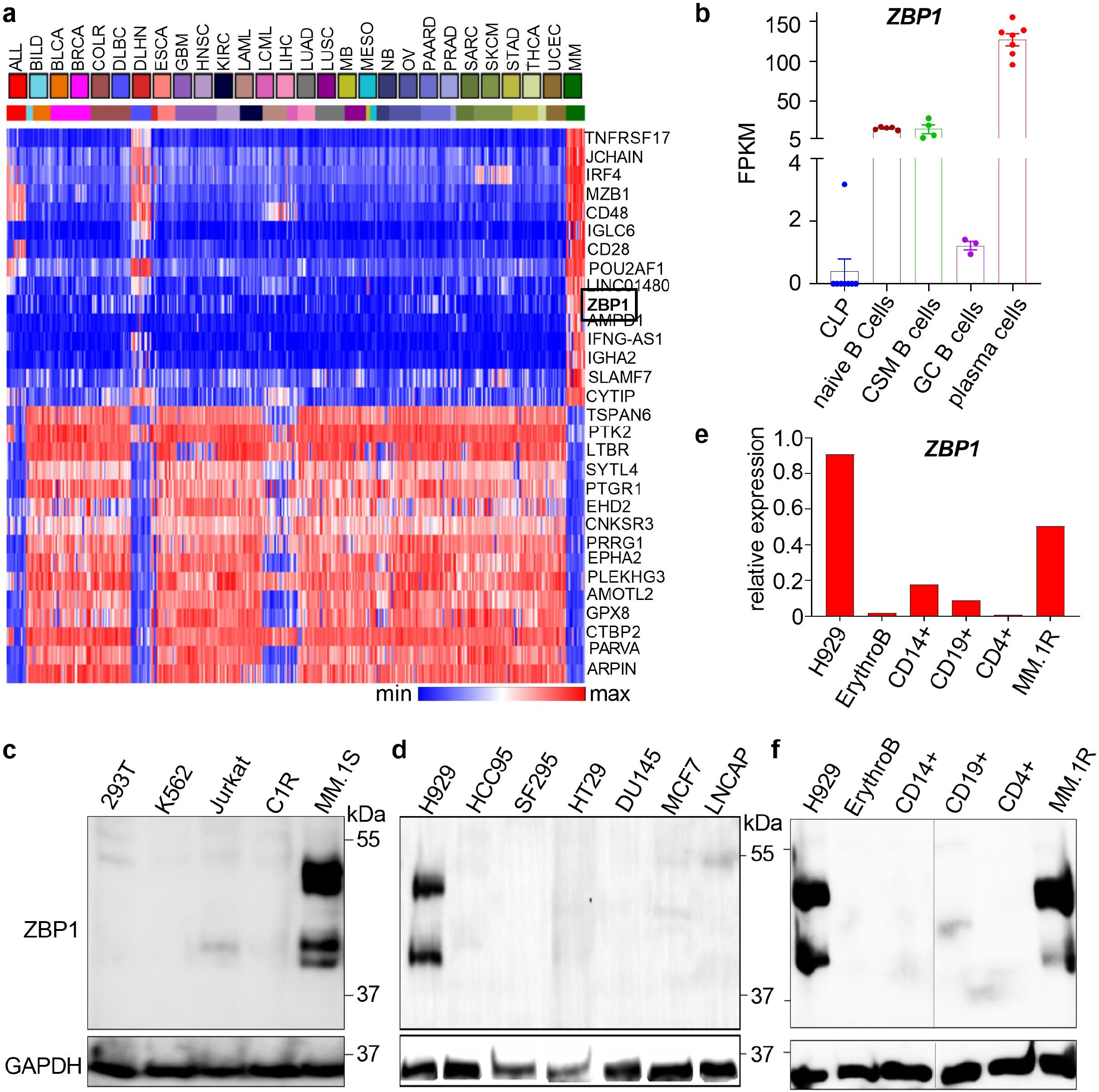
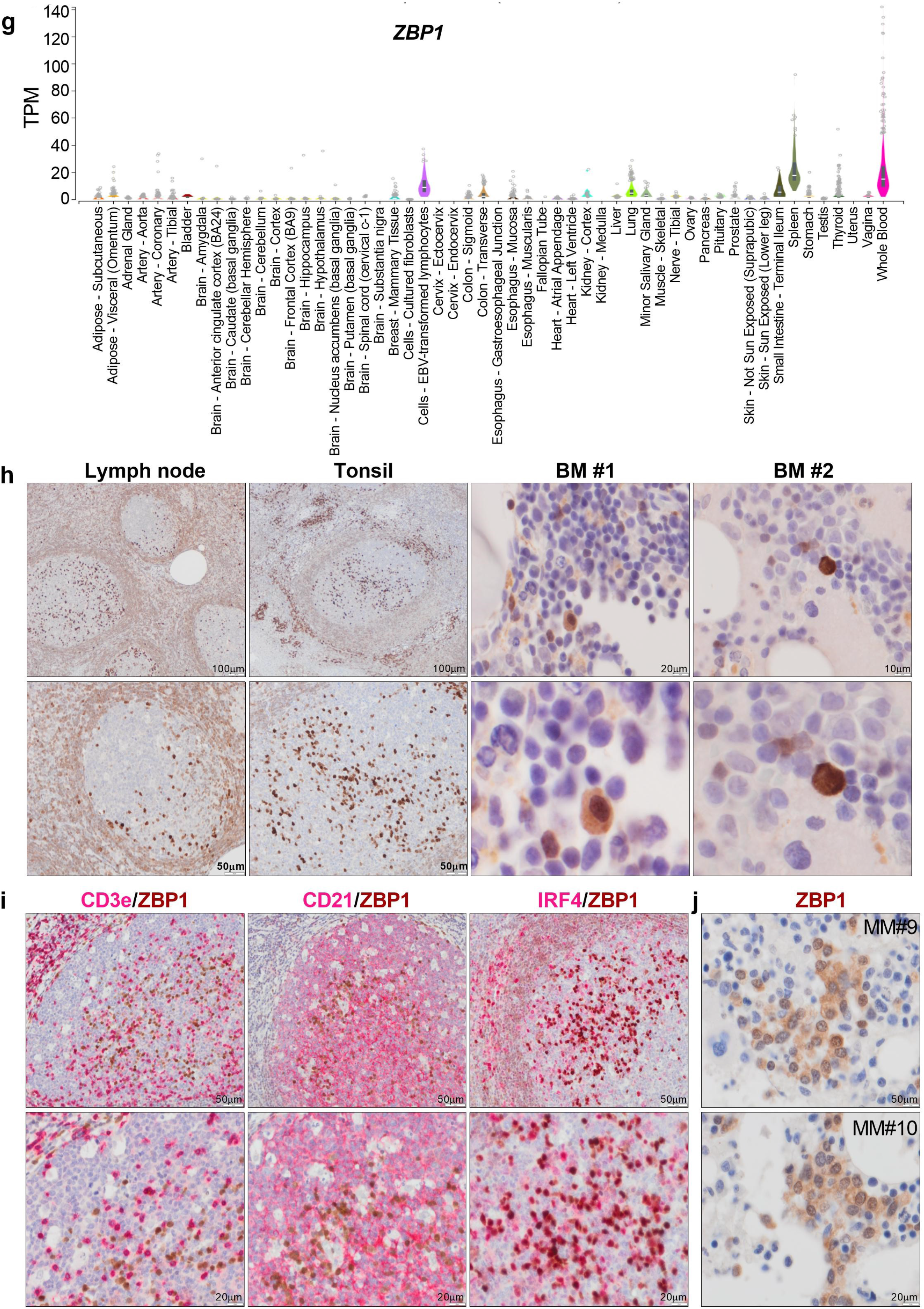
**(a)** Heat map showing differential expression of the top 30 genes between 27 MMCL and >800 other cancer cell lines (data from the Cancer Cell Line Encyclopedia Portal (CCLE)). *ZBP1* is boxed. ALL: Acute Lymphoblastic B Cell Leukaemia, BILD: Biliary Tract Carcinoma, BLCA: Bladder Carcinoma, BRCA: Breast Carcinoma, COLR: Colorectal Carcinoma, DLBC: Burkitt Lymphoma, DLHN: Lymphoma Hodgkin, ESCA: Squamous Cell Carcinoma, GBM: glioma HighGrade, HNSC: Upper Aerodigestive Tract, KIRC: Clear Cell Renal Cell Carcinoma, LAML: Acute Myeloid Leukaemia, LCML: Blast Phase Chronic Myeloid Leukaemia, LIHC: Hepatocellular Carcinoma, LUAD: Lung Non-Small Cell Carcinoma, LUSC: Lung Squamous Cell Carcinoma, MB: Medulloblastoma, MESO: Mesothelioma, NB: Neuroblastoma, OV: Ovary Carcinoma, PAARD: Pancreas Carcinoma, PRAD: Prostate Carcinoma, SARC: Ewings Sarcoma, SKCM: Skin Melanoma, STAD: Stomach Gastric Carcinoma, THCA: Thyroid Carcinoma, UCEC: Endometrium Carcinoma, MM: Plasma Cell Myeloma. **(b)** ZBP1 mRNA expression in human common lymphoid progenitors (CLP), naïve, class switched memory (CSM) and germinal centre (GC) B cells as well as tonsillar plasma cells (data from the Blueprint Portal; data shown as mean ± SEM). **(c, d)** Lack of ZBP1 expression in non-myeloma hematopoietic cell lines K562 (erythromyeloid), Jurkat (T cell lymphoblastic) and C1R (EBV-transformed B cell) cells and solid tumor cell lines HCC95 (squamous cell lung carcinoma), SF295 (glioblastoma), HT29 (colon adenocarcinoma), DU145 and LNCAP (prostate carcinoma), MCF7 (breast cancer). **(e, f)** ZBP1 mRNA and protein expression as assessed by qPCR (e) and immunoblotting (f) in purified primary human bone marrow erythroblasts, peripheral blood CD14+ monocytes, CD19+ B cells, CD4+ T cells. The MMCL H929 and MM.1S are shown as positive controls. **(g)** *ZBP1* mRNA expression in 53 human tissues. *ZBP1* expression is only detected in organs with lymphoid tissue (Data from the GTex portal; data shown as mean ± SEM). **(h)** ZBP1 expression in lymph nodes, tonsil, and healthy bone marrow as assessed by immunohistochemistry. Within lymph nodal and tonsillar germinal centres, ZBP1 expression is mostly restricted to PCs. Germinal centre B cells are mostly negative. ZBP1 is also strongly expressed in PCs outside the follicles. There is weaker ZBP1 expression in mantle cells and other interfollicular B cells. ZBP1 expression is mostly restricted to PCs in normal bone marrow (BM#1 & BM#2). **(i)** ZBP1 expression in tonsillar germinal centres co-stained with CD3e (T cells), CD21 (follicular dendritic cells) or MUM.1 (IRF4) as identified by immunohistochemistry. ZBP1 is not expressed in CD3e^+^ T cells or CD21^+^ follicular dendritic cells, but is co-expressed by IRF4 (MUM.1)^+^ PCs. **(j)** ZBP1 expression in bone marrow myeloma PC from another two patients with MM.

**Supplementary Fig. 2 (Related to Fig. 3):**
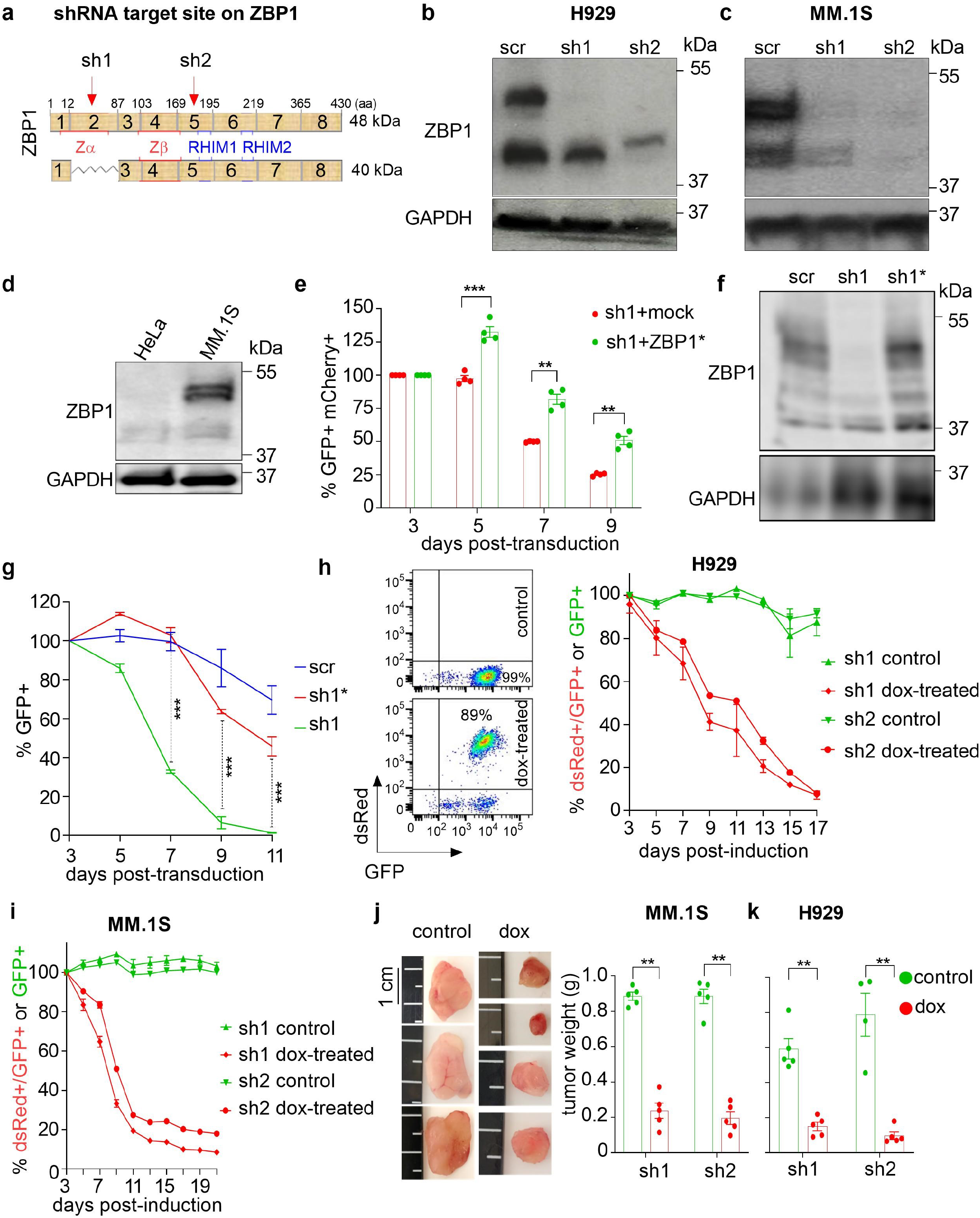
**(a)** Schematic of *ZBP1* mRNA exonic structure, positions of Zα, Zβ and RHIM domains and of shRNA1 and shRNA2 with reference to the two main ~48 and ~40 kDa ZBP1 isoforms are shown. **(b, c)** Immunoblotting against ZBP1 in H929 and MM.1S MMCL 4 days after lentiviral transduction with sh1 or sh2 or scr control. GAPDH is shown as loading control (n=3). **(d)** Immunoblotting of ZBP1 expression in the epithelial cancer cell line HeLa. **(e)** Percent GFP^+^ cells after co-transduction of MM.1S myeloma cells with *ZBP1*-targeting sh1 plus empty vector (mock) or with *ZBP1*-targeting sh1 plus *ZBP1* cDNA with mutated seed annealing sequences (ZBP1*) (n=4). **(f)** ZBP1 expression assessed by immunoblotting in MM.1S cells transduced with shRNA1 (sh1), ‘seed’shRNA1 (sh1*) or scr control. **(g)** Percent GFP^+^ cells after transduction of MM.1S myeloma cells with ZBP1-targeting sh1, ‘seed’ shRNA1 (sh1*) or scr control (n=3). **(h, i)** Representative flow-cytometric analysis of doxycycline inducible shRNA1 in MMCL before and after dox treatment (left). Transduced cells are GFP+, cells with dox-induced shRNA are dsRed+. Percent dsRed+ H929 (h) and MM.1S (i) myeloma cells with and without dox treatment *in vitro* (right) (n=3). **(j, k)** Photographs of tumors explanted at sacrifice from control (i.e., non-dox-treated) or dox-treated animals engrafted with MMCL MM.1S transduced with dox-inducible shRNA1&2 targeting ZBP1 and tumor weight at sacrifice in animals engrafted with MM.1S (j) andH929 (k) myeloma cells transduced with anti-ZBP1 shRNA or scr control. Data shown as mean ± SEM, unpaired t-test; ** p≤ 0.01, *** p≤0.001.

**Supplementary Fig. 3 (Related to Fig. 3):**
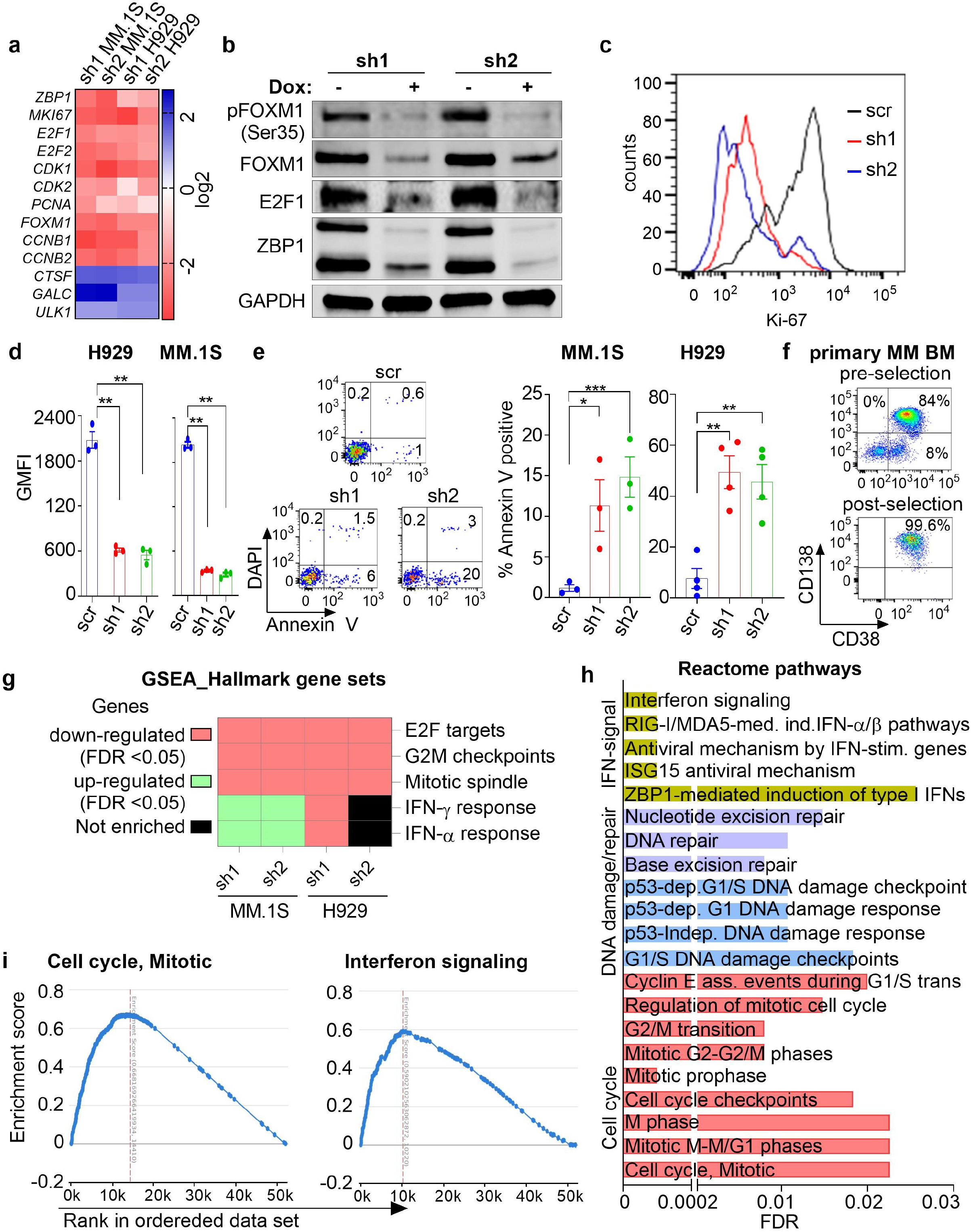
**(a)** Heatmap showing log_2_-fold change in expression of genes of interest 4 days after transduction of MMCL with anti-ZBP1 sh1 or sh2 or scr control (p adj <0.05). **(b)** Immunoblotting of genes of interest on day 3 following Dox-induction of anti-ZBP1 sh1 or sh2 in MM.1S myeloma cells. **(c, d)** Flow-cytometric analysis of Ki-67 expression in MM.1S myeloma cells (c) and cumulative data in H929 and MM.1S MMCL (d) (n=3). **(e)** Flow-cytometric analysis of apoptosis by Annexin V staining and cumulative data in MM.1S and H929 MMCL (n=3). **(f)** Representative flow-cytometric example of bone marrow-derived myeloma PC purity before and after CD138 immunomagnetic bead selection. **(g)** Gene set enrichment analysis with reference to Molecular Signatures Database v7.1 Hallmark gene sets of transcriptomes of ZBP1-depleted MM.1S and H929 MMCL. **(h)** GSEA of highest 5% vs lowest 90% *ZBP1-*expressing myeloma PC. Analysis was performed using RNA-seq transcriptomes of purified myeloma PC (n=767 patients) and the GSEA tool inbuilt in the multiple myeloma research foundation (MMRF) research gateway portal. **(i)** GSEA example of reactome pathway for cell cycle, mitotic genes enriched in the top 5% *ZBP1*-expressing cohort. Error bars represent mean ± SEM, unpaired t-test; ** p≤ 0.01, *** p≤0.001.

**Supplementary Fig. 4 (Related to Fig. 4):**
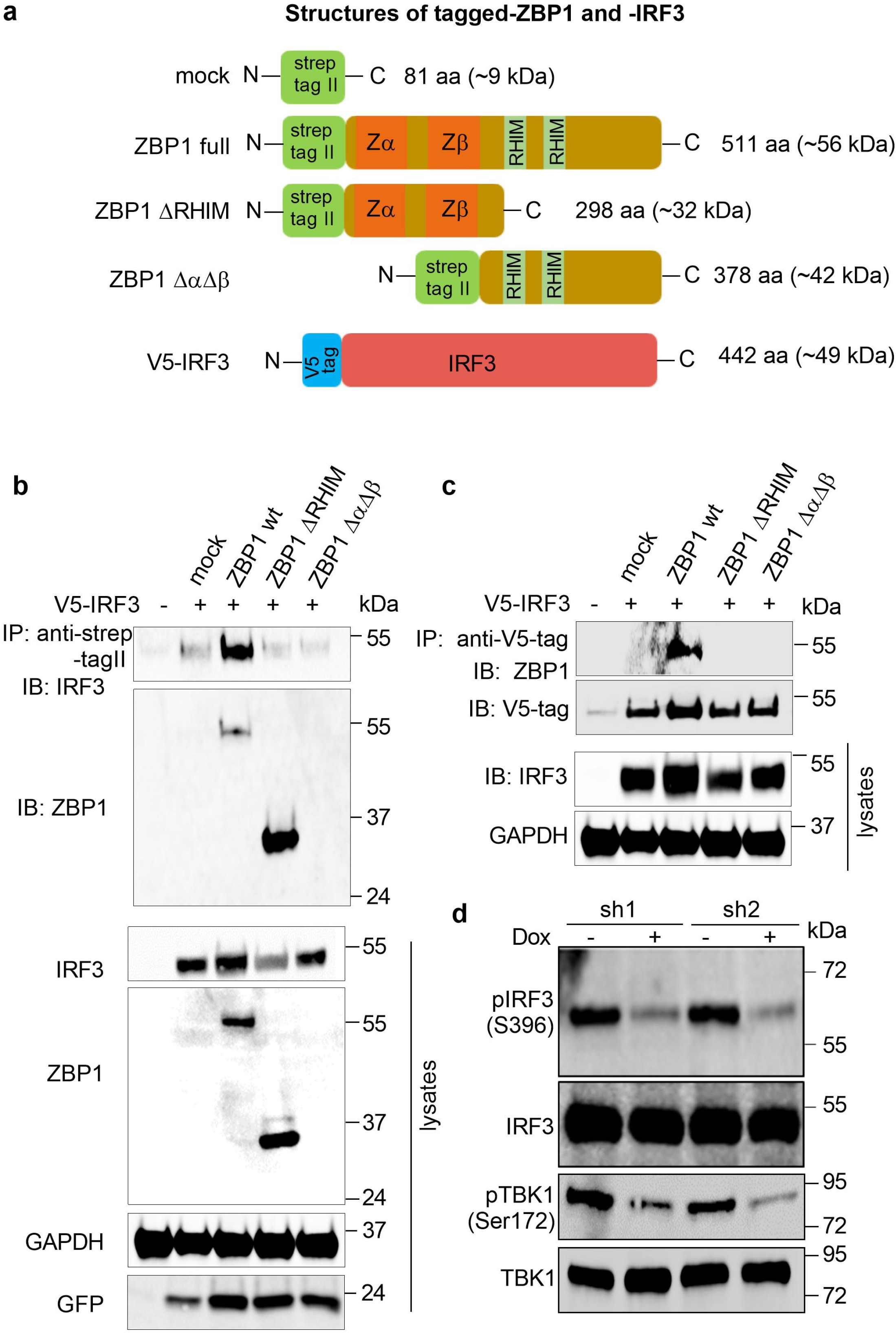
**(a)** Schematic illustration of V5-tagged IRF3 and strep-tag II full length ZBP1 and its deletion mutants. RHIM, rip homotypic interaction domain. **(b, c)** Strep-tagII ZBP1-full, ZBP1-ΔRHIM, ZBP1-ΔαΔβ and mock GFP-expressing cDNA constructs, were transiently co-expressed with V5-tagged IRF3 in HEK293T cells. Cell lysates were immunoprecipitated with anti-V5 antibody or anti-strep-tag magnetic beads followed by western blotting with anti-ZBP1 or -IRF3 antibody. IRF3 can be readily co-immunoprecipitated with only full length ZBP1. Note that anti-ZBP1 Ab used here binds an N-terminal epitope that is removed in the ZBP1-ΔαΔβ construct and therefore expression of transfected ZBP1-ΔαΔβ cannot be detected directly. Similarly, anti-StrepTag Ab does not work in immunoblotting although it is efficient in immunoprecipitation. **(d)** Immunoblot against pIRF3/IRF3 and pTBK1/TBK1 in dox-induced or non-induced ZBP1-depleted cells.

**Supplementary Fig. 5 (Related to Fig. 4):**
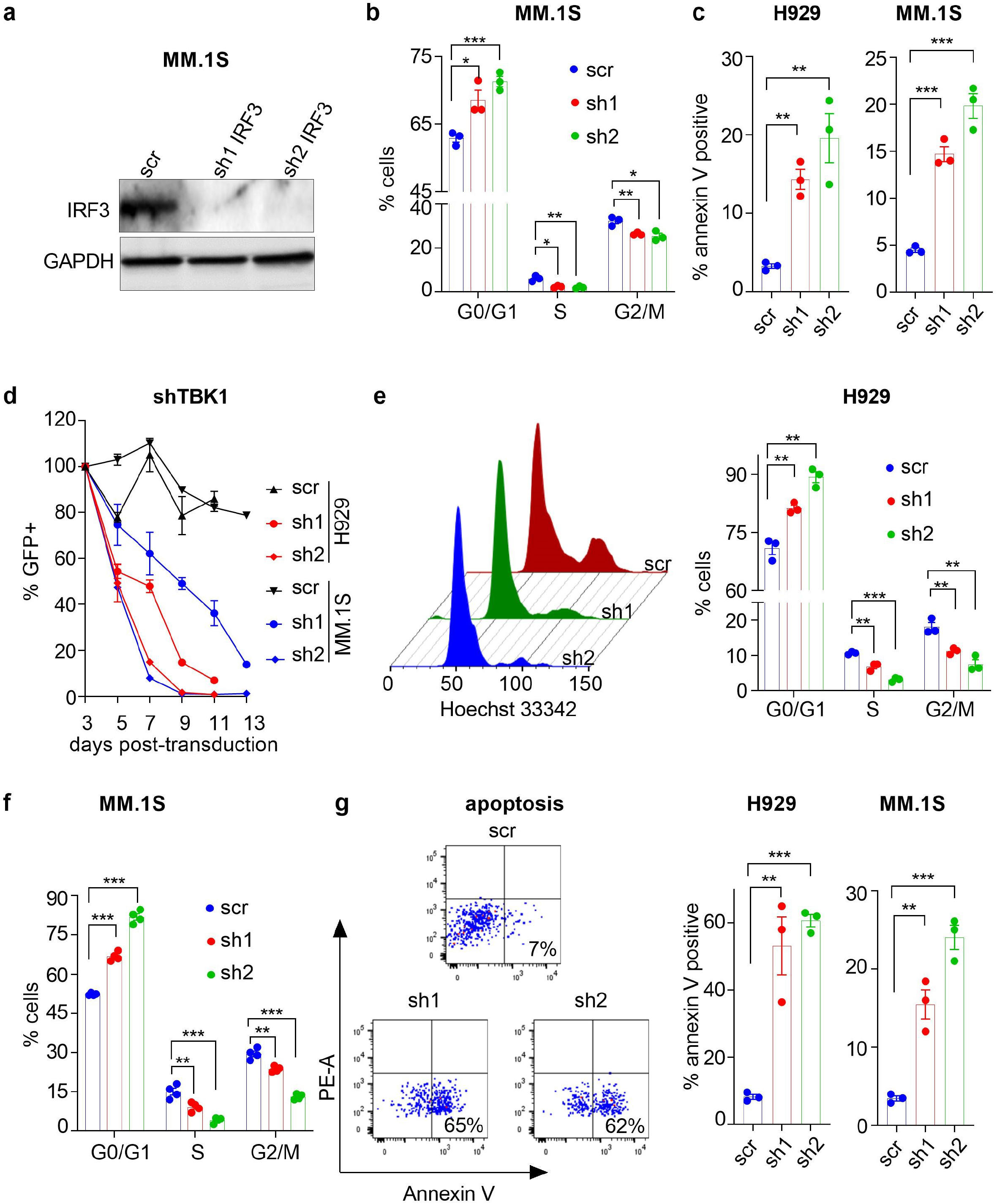
**(a)** Immunoblot against IRF3 in MM.1S cells 4 days post-transduction with scr or IRF3 shRNA1 or shRNA2. **(b)** Flow-cytometric analysis of cell cycle in MM.1S cells 4 days post-transduction with scr or IRF3 shRNA1 or sh2 (n=3). **(c)** Annexin V staining for assessment of apoptosis in H929 and MM.1S cells on day 4 post-transduction with scr or IRF3 shRNA1 or shRNA2 (n=3). **(d)** Percent GFP^+^ cells after transduction with scr control or shRNA1 or shRNA2targeting TBK1 in H929 and MM.1S cells. Normalised to day 3 GFP expression for each shRNA or scr control (n=3). **(e, f)** Flow-cytometric analysis of cell cycle in H929 (e) (n=3) and MM.1S (f) cells on day 4 post-transduction with *TBK1*-depleting shRNA1 or shRNA2 or scr (n=4). **(g)** Annexin V staining for apoptosis in MMCL H929 and MM.1S cells on day 4 post-transduction with *TBK1*-depleting shRNA1 or shRNA2or scr control (n=3). Data shown as mean ± SEM, unpaired t-test; * p≤ 0.05, ** p≤ 0.01, *** p≤0.001.

**Supplementary Fig. 6 (Related to Fig. 5):**
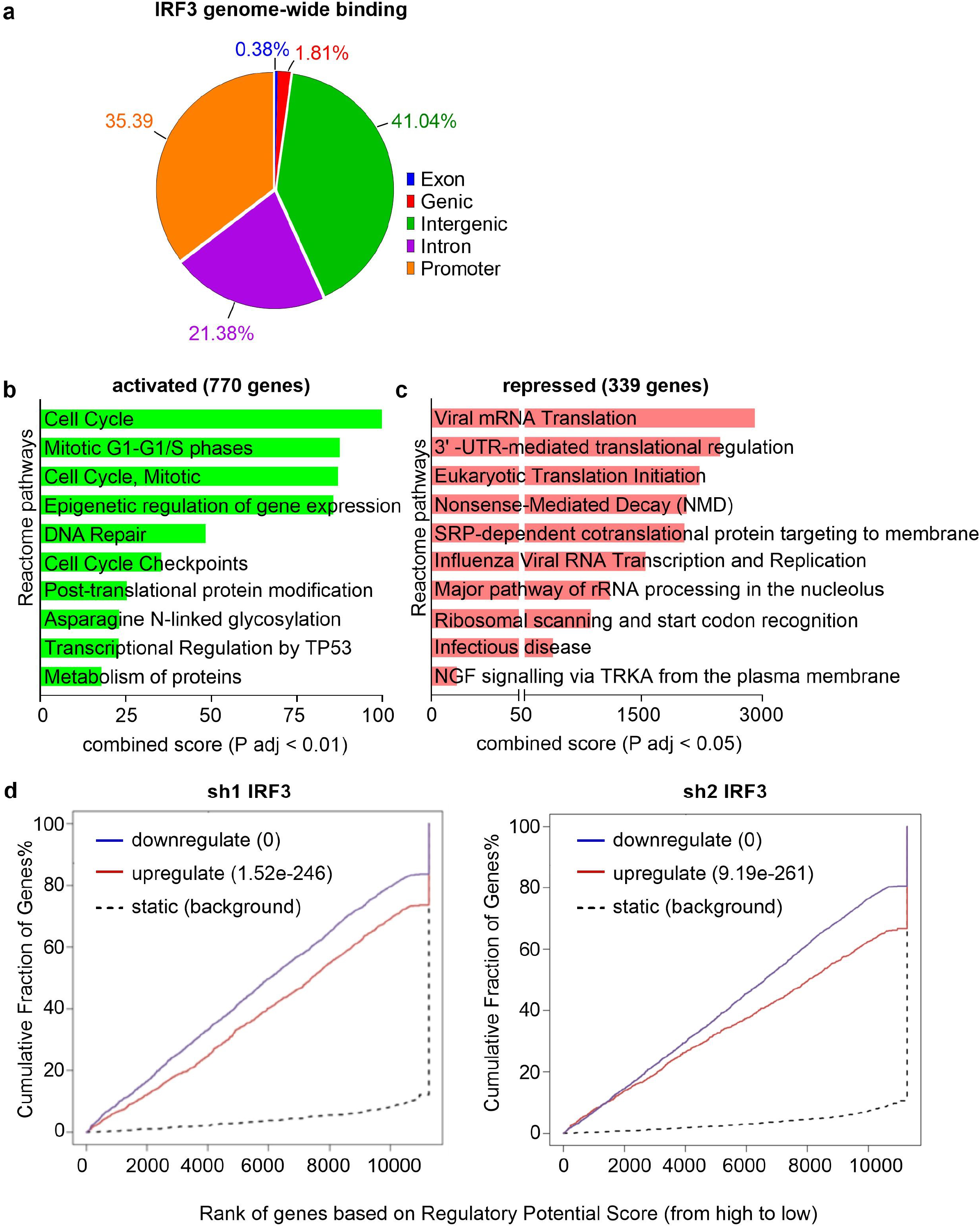
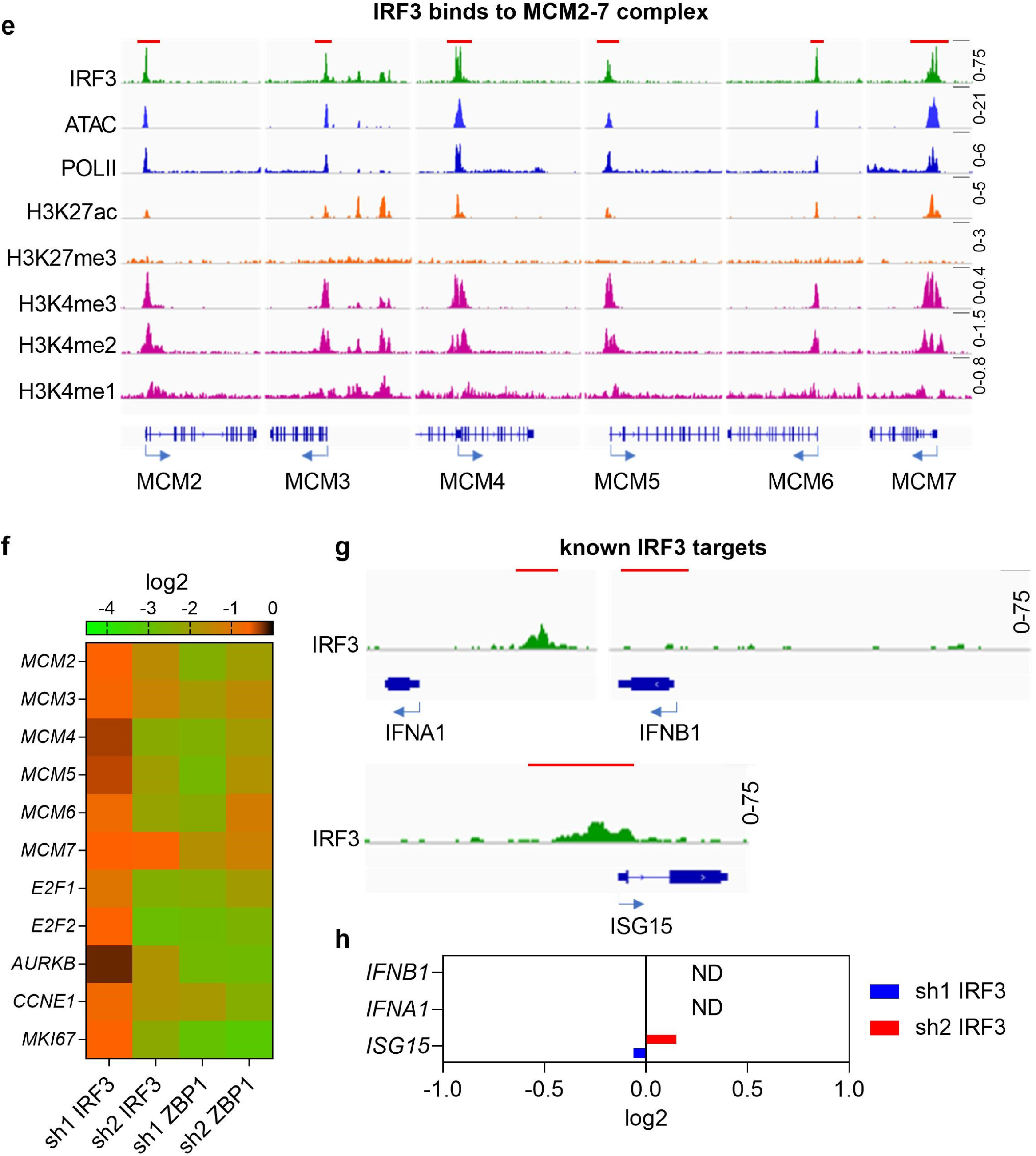
**(a)** Annotation of IRF3 genome wide binding according to genomic features. **(b, c)** Enrichr pathway enrichment analysis of the 770 and 339 genes predicted to be directly activated (b) or repressed (c) by IRF3 binding to their regulatory areas. **(d)** Regulatory potential prediction models display significant activating (blue) and repressive (red) function of IRF3 in MM.1S cells. Models derived from BETA-plus analysis, after integrating the IRF3 cistrome with *IRF3*-depleted transcriptome obtained by RNA-seq analysis for shRNA1 or shRNA2. **(e)** IGV browser snapshots of IRF3 and Pol II binding, chromatin accessibility and histone mark enrichment at regulatory areas of several genes promoting cell cycle progression and cell proliferation. The red block on the top indicates 5kb. **(f)** Heatmap of the indicated mRNA expression levels in indicated mRNA-depleted RNA-seq data (p adj <0.05). **(g, h)** IRF3 binding to IFN type I pathway genes (g), the red block on the top indicates 1kb, and mRNA expression change of indicated genes (h). Note neither *IFNA1* nor *IFNB1* are expressed before or after IRF3 depletion.

**Supplementary Fig. 7 (Related to Fig. 6):**
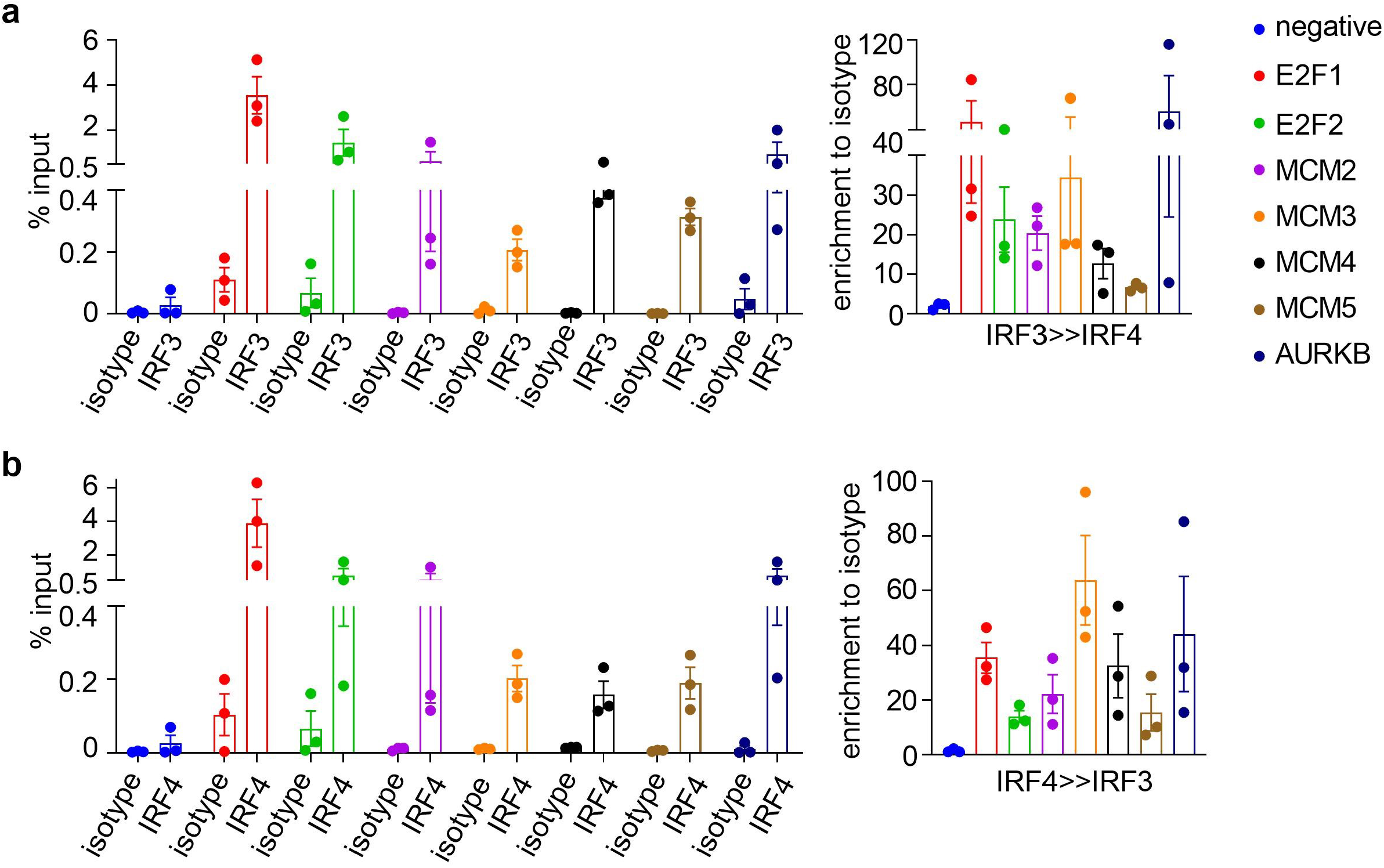
**(a)** qPCR of IRF3 ChlP or IRF3 to IRF4 ChlP-re-ChlP (n=3). **(b)** qPCR of IRF4 ChlP or IRF4 to IRF3 ChlP-re-ChlP (n=3).

## Notes

### Competing Interest Statement

The authors have declared no competing interest.

